# Short-chain di-carboxylates as positive allosteric modulators of the pH-dependent pentameric ligand-gated ion channel GLIC: requirement of an intact vestibular pocket

**DOI:** 10.1101/2023.03.03.530963

**Authors:** Catherine Van Renterghem, Ákos Nemecz, Sandrine Delarue-Cochin, Delphine Joseph, Pierre-Jean Corringer

**Affiliations:** Institut Pasteur, Université de Paris, CNRS UMR 3571, Channel-Receptors Unit, F-75015 Paris, France; Université Paris-Saclay, CNRS, BioCIS, F-92290, Châtenay-Malabry, France

**Keywords:** pLGIC, GLIC, fumarate, succinate, positive allosteric modulator, orthosteric site, orthotopic site, vestibular site, intracellular pH.

## Abstract

GLIC is a prokaryotic orthologue of brain pentameric neurotransmitter receptors. Using whole-cell patch-clamp electrophysiology in a host cell line, we show that short-chain di-carboxylate compounds are positive modulators of pHo 5-evoked GLIC activity, with a rank order of action fumarate > succinate > malonate > glutarate. Potentiation by fumarate depends on intracellular pH, mainly as a result of a strong decrease of the pHo 5-evoked current when intracellular pH decreases. The modulating effect of fumarate also depends on extracellular pH, as fumarate is a weak inhibitor at pHo 6 and shows no agonist action at neutral pHo. A mutational analysis of residue-dependency for succinate and fumarate effects, based on two carboxylate-binding pockets previously identified by crystallography (Fourati *et al*. 2020), shows that positive modulation involves both the inter-subunit pocket, homologous to the neurotransmitter-binding orthotopic site, and the intra-subunit (also called vestibular) pocket. An almost similar pattern of mutational impact is observed for the effect of caffeate, a known negative modulator. We propose, for both di-carboxylate compounds and caffeate, a model where the inter-subunit pocket is the actual binding site, and the region corresponding to the vestibular pocket is required either for inter-subunit binding itself, or for binding-to-gating coupling during the allosteric transitions involved in pore gating modulation.

**Key points summary:**

- Using a bacterial orthologue of brain pentameric neurotransmitter receptors, we show that the orthotopic/orthosteric agonist site and the adjacent vestibular region are functionally inter-dependent in mediating compound-elicited modulation. We propose that the two sites in the extracellular domain are involved “in series”, a mechanism which may have relevance to Eukaryote receptors.
- We show that short-chain di-carboxylate compounds are positive modulators of GLIC. The most potent compound identified is fumarate, known to occupy the orthotopic/orthosteric site in previously published crystal structures.
- We show that intracellular pH modulates GLIC allosteric transitions, as previously known for extracellular pH.
- We report a caesium to sodium permeability *ratio* (P_Cs_/P_Na_) of 0.54 for GLIC ion pore.

## Introduction

The use of Prokaryote pentameric ligand-gated ion channels (pLGICs; or Pro-Loop receptors, Jaiteh *et al*. 2016) as models for brain pentameric neurotransmitter receptors was developed for a pragmatic reason: it is less difficult to purify and crystallize a prokaryotic than a mammalian membrane protein. And indeed two Prokaryote receptors, ELIC and GLIC, provided the first crystal structures in the pLGIC group of proteins (Hilf & Dutzler 2008, 2009; Boquet *et al*. 2009). GLIC provided a series of different conformational states (Prevost *et al*. 2012), and the first model at atomic resolution for the transition between a closed state and a (most probably) open state (Sauguet *et al*. 2014). GLIC and ELIC appear as excellent models for mammalian pLGICs, since the main structural, biophysical and pharmacological properties are very well conserved.

But Prokaryote pLGICs also present another interest regarding brain receptors studies: would new questions originate from the few differences occurring? Compared anatomy, studying animal species at various stages of evolution, brought a concept such as metamery, which, together with embryology, allowed a much more conceptualized view of the human body anatomy. Although the receptor channels GLIC and ELIC are much more similar to brain pLGICs than a fish or a bird is similar to a *Homo sapiens*, differences occurring in Prokaryote pLGICs may lead to question concepts developed in the study of mammalian receptors, and re-contextualize them in a wider view. Would previously non considered points regarding human pLGICs emerge from prokaryotic studies?

A vestibular, intra-subunit compound binding pocket (one per subunit) was identified from X-ray crystallography in the extracellular domains (ECDs) of three Prokaryote pLGICs, ELIC (Spurny *et al*. 2012), GLIC (Nury *et al*. 2010, Sauguet *et al*. 2013, Fourati *et al*. 2015, 2020), and sTeLIC (Hu *et al*. 2018). Such a vestibular binding site is not known in mammalian pLGICs: the vestibular binding site was not selected during evolution, leaving full place to the orthotopic site, as neurotransmitter binding site (in this report, we use the word “orthotopic” rather than the more usual word “orthosteric” to designate the pLGICs reference binding site, see Methods). What is remaining from this vestibular site in neurotransmitter receptors? Is it possible to “awake” a functional vestibular drug binding site in human pLGICs (this would open the way to a new class of pharmacological agents)?

The last point derives from the assumption that the vestibular pocket is a functional binding site in Prokaryote pLGICs: is this point established? Is the vestibular pocket functional as an agonist binding site? This hypothesis was proposed in the case of sTeLIC, a high extracellular pH (high pHo) activated pLGIC, for the quasi-agonist action of 4-bromocinnamate, on the basis that 4-bromocinnamate was identified in the intra-subunit pocket, in the available co-structure, while the orthotopic site was empty (Hu *et al*. 2018).

Is the vestibular pocket functional as an (allotopic) binding site for a positive or negative modulator of the agonist action of a compound binding to the orthotopic site? This is the conclusion proposed in the case of ELIC, in the report which first emphasized a vestibular binding pocket in a pLGIC (Spurny *et al*. 2012). The vestibular pocket (five per homopentamer) was identified by crystallography, as a benzodiazepine (flurazepam) binding pocket, while bromoflurazepam occupied the inter-subunit sites which were not occupied by the ELIC agonist GABA. On ELIC activated by GABA, flurazepam was shown to produce a potentiating effect at low concentration, and an inhibitory effect at high concentration. The authors attributed the potentiating action to vestibular site binding, on the basis that two mutations (ELIC N60C and I63C) in the vestibular/intra-subunit pocket suppressed the potentiating effect. They attributed the inhibitory action to orthotopic site binding, on the basis that a mutation in the orthotopic site (F19A, corresponding to Y23A in GLIC) suppressed the benzodiazepine inhibitory effect (ELIC Phe19 was chosen because it is in contact with flurazepam, not with GABA). The authors conclude that the intra-subunit site mediates potentiation and the inter-subunit site mediates inhibition.

In the present report, we show that the vestibular pocket is functional as a “coupling” region, whose integrity conditions the efficiency (or even the binding) of the orthotopic site ligand. In GLIC, two carboxylate-binding pockets were previously characterized by crystallography (Sauguet *et al*. 2013, Fourati *et al*. 2015, 2020): 1/ an inter-subunit (inter-SU) pocket, overlapping with the location homologous to the conserved orthotopic agonist site, but slightly more deeply buried, as the GABA binding pocket in ELIC, and 2/ an intra-SU pocket, exactly superimposed to the vestibular pocket of ELIC.

We show that short-chain dicarboxylic acid/carboxylate (di-CBX) are positive modulators of the allosteric transitions (PAMs) in GLIC activated at low extracellular pH (pHo). Fumaric acid/fumarate (FUMAR) is the best PAM identified. We show that a ditetrazole fumarate bioisostere mimics FUMAR effect, whereas other fumarate structural analogs are inactive.

Using seven GLIC variants with a single mutation in one of the CBX-binding pockets, we show that the di-CBX PAM effect is fully abolished by single mutations in the conserved orthotopic/inter-SU pocket, but also by single mutations in the vestibular pocket. A similar “all-or-none” pattern of mutational impact (in both orthotopic/inter-SU *and* vestibular pockets) is observed for the negative modulator of allosteric transitions (NAM) effect of caffeic acid/caffeate (CAFFE). On the basis of this “all-or-none” pattern of mutational impact, and published crystallographic data, we propose a “in series” mechanism, in which binding occurs at the orthotopic site, as in Eukaryote pLGICs, and the vestibular region is involved in, and conditions, binding-to-gating coupling (or may condition orthotopic binding itself). The “in series” mechanism is valid for both positive (di-CBX) and negative (CAFFE) modulation.

In the associated report (Van Renterghem *et al*., Manuscript M2), we establish a different, “loose” pattern of impact of the same CBX-pockets single mutations, on mono-CBXs NAM effects. From the “loose” pattern of mutational impact, and from the properties of a pre-β5 double mutant, while keeping the “in series” mechanism, we propose that, in addition to being a coupling region, the vestibular pocket may function as an accessory (allotopic) binding site, exerting a *veto* in-between the main orthotopic binding site and the gating machinery. The two reports question the relevance of a vestibular pocket regarding human pLGIC modulation, a point also addressed by Brams *et al*. (2020).

Additionally, we report here a permeability *ratio* P_Cs_/P_Na_ near 0.54 for GLIC at pHo 5. We show that, unexpectedly, the di-CBX PAM effect occurs only in the upper part of the activation curve (by extracellular protons). Finally, using a low-pH pipette solution, we show that the value of GLIC current activated at pHo 5.0 becomes smaller when intracellular pH (pHi) decreases, while the current activated by FUMAR at pHo 5.0 is much less reduced, leading to a much larger FUMAR potentiation *ratio* at low pHi. We discuss this data in terms of 1/ GLIC allosteric modulation by pHi, 2/ an explanation for the great variability of GLIC electrophysiology data, and 3/ GLIC desensitization *vs* pHi-controlled GLIC current decay.

## Methods

### Host cell line

Whole-cell recordings showed that HEK293 and CHO cells were able to express GLIC after transfection. However, they both showed contaminating low-pHo activated currents, arising from endogenous channels. Gunthorpe *et al*. (2001) reported the presence of acid-sensing ion channels (ASICs), specifically of ASIC-1 type, in HEK293 cells. In recordings from non-transfected HEK293 cells [cultured in Dulbecco’s Modified Eagle Medium with Glutamax, with 10 % heat-inactivated fetal bovine serum (dFBS), penicillin (0.1 g/L), and streptomycin (100 U/mL)], we observed 1/ a fast, transient low-pHo induced increase in a conductance selective for sodium ions, with a peak value reaching 80% of maximum near pH 6, and 2/ a delayed, sustained increase in a conductance selective for multiple cations over chloride anion, with a plateau value still increasing between pH 5 and pH 4. These properties are compatible with those of ASIC-1 (Waldmann *et al*. 1997) and non-type-1 (Waldmann & Lazdunski 1998) ASIC channels respectively. Recordings from non-transfected CHO cells [Ham’s F12 medium with 5% dFBS, 2 mM glutamine and the same antibiotics] revealed ASIC-1 type currents. Such proton-activated ASIC type currents were absent in recordings from non-transfected tk-ts13 cells, from a thymidine-kinase negative (Waechter & Baserga 1982), temperature-sensitive variant (Talavera & Basilico 1977) of the Baby Hamster Kidney (BHK) cell line. Therefore, the tk-ts13 cell line was selected as a host for the patch-clamp study of GLIC.

### GLIC expression in tk-ts13 cells

Cells were grown under a 5% CO_2_ atmosphere at 37°C (an upper-limit value for these cells which do not grow at 39°C), in Dulbecco’s Modified Eagle Medium with *Glutamax*, glucose and pyruvate (Invitrogen), supplemented with dFBS (5 %), and penicillin (0.1 g/L) plus streptomycin (100 U/mL), or no antibiotics in late experiments. Cells in Petri dishes (35 mm in diameter) were transfected one day after cell seeding (2-10 x 10^4^ cells along a “+” in the middle part of the dish), using a calcium phosphate-DNA co-precipitation method. A pMT3 plasmid containing the DNA sequence of GLIC was mixed with a pMT3 plasmid coding for GFP, in quantities of 2 and 0.2 µg DNA per 35 mm dish. Isolated, GFP-positive cells were used for electrophysiology one or two days after transfection, as there was little or no current on GFP-positive cells recorded on the third day. GLIC residue numbering follows PDB entry 3EAM (Bocquet *et al*. 2009), in which number 5 was assigned to the (N-terminal) Val4 of GLIC, so that Arg77 in the present report corresponds to Arg76 GLIC residue.

### Whole-cell patch-clamp electrophysiology

Unless otherwise specified (*i*.*e*. in Fig. 6, from oocytes), the data presented was obtained from GFP-positive tk-ts13 cells voltage-clamped in the whole-cell configuration. Patch-clamp methods (Hamill *et al*. 1981) were up-dated from Van Renterghem & Lazdunski (1991, 1994). Whole-cell voltage-clamp currents were recorded using an RK-400 patch-clamp amplifier (Bio-Logic, France; discontinued) with a HK-R-410/08 dual resistor headstage, connected to a computer using a Digidata 1550 analog/digital – digital/analog converter, and the program pClamp 10 (Axon Instruments). Currents were low-pass filtered (RK-400 internal 5-poles Bessel filter) at 1 kHz, and digitized at a sampling frequency of 5-10 kHz. Digital filtering by 500-10000 to 1 data points averaging was further applied to display traces in the Figures. Pipettes were pulled from thick wall (0.37 mm) borosilicate glass (1.5 mm o.d., 0.75 mm i.d.), and fire-polished to resistances of 2.8 to 4.5 MΩ in our standard solutions. The standard pipette solution was composed of (in mmol/L (mM)): CsCl (150), MgCl_2_ (1), BAPTA (10), HEPES (10), and CsOH to pH 7.3. The standard extracellular solution was composed of (in mM): NaCl (165), MgCl_2_ (1), CaCl_2_ (1), MES (10) and HEPES (6), pH 7.5 (NaOH to 9.5 / HCl). In most cases, solutions were applied locally near the recorded cell, using a gravity driven, multiway perfusion system, converging to a single tip (50-100 µL/min, exchange time # 1 s), with or without computer-driven valves. Most recent experiments (Figs. 3, 5*C-F*) used a motorized head, rapid solution exchange system (RSC-200, Bio-Logic), ending with parallel 1 mm o.d. glass tubes. Exchange time (as judged from the open-tip pipet / diluted solution technic) was <30 ms under our conditions (using a light holding micro-manipulator). An agar bridge (a water, KCl 3 M, agar 5 g/L gel, in a bended piece of thin-wall patch pipet glass tube) was used to isolate the extracellular reference electrode from changes in pH (a useful precaution in case any silver oxide would contaminate the silver/silver chloride pellet electrode). For the intracellular, pipette electrode, a silver wire was covered with AgCl by electrolytic oxidation of Ag into Ag^+^ in a 0.1 M KCl water solution (against a platinum electrode). Electric current flowing through the membrane from extracellular to intracellular face is counted negative, and represented downwards in the Figures.

### GLIC expression in *Xenopus* oocytes, and oocyte electrophysiology

GLIC mutants were characterized for any gain- or loss of function sensitivity to extracellular protons (Fig. 6), using whole-oocyte two-electrode voltage-clamp current recording (Kusano *et al*. 1982) after receptor-channel membrane expression (Sumikawa *et* a*l*. 1981). GLIC expression (nuclear injection of GLIC (0.08 g/L) and GFP (0.02 g/L) cDNAs in separate pMT3 vectors), and recording conditions, were as in Nemecz *et al*. 2017, except that a motorized, fast chamber-perfusion system allowed fast solution exchange in a small-volume (0.5 mL) recording chamber (ÁN). A GeneClamp500 (Axon Instruments; discontinued) voltage-clamp amplifier was used, with two intracellular electrodes connected to the cytosol *via* two 3 M KCl filled glass pipettes (0.2-1.0 MΩ), and two amplifier virtual ground electrodes connected to the bath *via* two 3 M KCl-agar bridges. The standard extracellular solution contained (mM) NaCl (100), KCl (3), MgCl_2_ (1), CaCl_2_ (1), MES (10), pH 8.0 (NaOH to pH 9.5 / HCl).

### Preparation of solutions

As GLIC is cation-selective, all solutions were prepared from the acid forms of the buffers/acido-basic compounds (BAPTA, HEPES, MES, di-CBX, etc.). Extracellular solutions were prepared as 10x stocks kept at 4 °C for a month. Full neutralization, up to pH 9.5 (NaOH), was followed by return (HCl) to pH 7.8 or 8.3 (for patch and oocyte), before volume adjustment. These 10x stocks were diluted to prepare the standard extracellular solutions, at pH 7.5 (patch) or 8.0 (oocyte). Lower pH values were reached by adding HCl 2M in water, and more pH 7.5/pH 8.0 solution if required. The patch pipette solution was prepared 1x, with partial neutralization (CsOH) up to pH 7.3. It was aliquoted and kept at -20 °C for a year. The lower pH value (6.0) was reached by adding a 2M HCl water solution.

### Counter-cation in di-CBX solutions

Effective di-CBX compound concentrations are up to 20 mM for GLIC, and the channel is cation selective. Following neutralization of carboxylic acids, the *ratio* ρ of added monovalent cation concentration to CBX concentration at pH 5.0 is between 0.6 and 0.7 for mono-, and between 1.1 and 2.0 for the di-CBXs considered (see Table 1). We found that adding 30 mM NaCl to our NaCl-based extracellular solution increased pHo 5.0-induced GLIC inward current at -40 mV by about 15 %. This increase in current was probably due to the increase in the GLIC-permeant cation concentration outside the cell (here Na^+^), as predicted in the constant field model by the voltage and current equations (Goldman 1943; Hodgkin & Katz 1949), rather than to a modulation of GLIC gating by Na^+^ ions. This extra-sodium induced increase in current was considered to be a problem.

The obvious protocol was to neutralize the carboxylic acids with a base giving a non-permeant cation. And indeed, the six-carbon cation resulting from the neutralization of *N*-methyl-D-glucamine (NMDG) is widely used as a non-permeant cation substituent in electrophysiological studies. However, whole-cell recordings showed that 10 mM succinic acid neutralized by NMDG base, as well as 20 mM NMDG base neutralized by HCl, both produced a slowly developing, complete and reversible inhibition of GLIC current at pHo 5.0. This inhibitory effect on GLIC has to be attributed to the NMDG^+^ cation, 4.5 pH-units under the NMDG^+^/NMDG p*K*a value (# 9.5). The inhibitory effect on GLIC showed no obvious voltage-dependent variation, it is therefore probably not due to an open channel block mechanism.

We therefore went back to a NaOH neutralization of carboxylic acids. But in the case of di-CBXs, we chose to prepare solutions with a *constant total Na^+^ ion concentration* by reducing the amount of NaCl provided. This in turn produced di-CBX solutions with a deficit in the Cl^-^ concentration, and a slightly reduced osmotic pressure, which all seemed preferable to an increased Na^+^ concentration. Attention to this detail excluded any contribution of increased Na^+^ concentrations to the di-CBX-induced increase in GLIC current reported here.

**Table 1.**
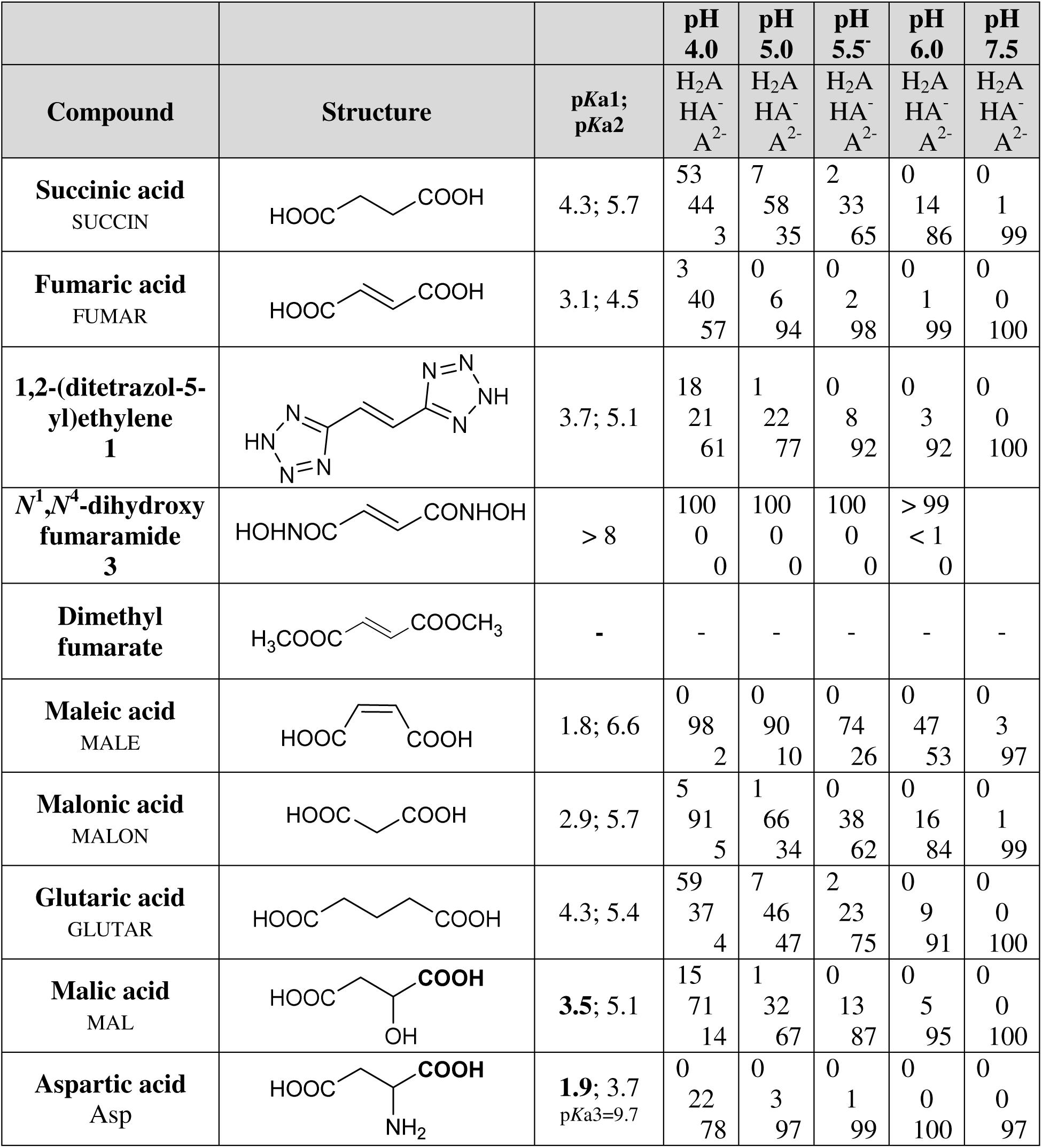
Di-CBX compounds and FUMAR analogs: structure, acidity constants and species distribution according to pH. Acidity constants *K*a1 and *K*a2 (given as p*K*a=-log(*K*a)) and species distributions were obtained from the program Dozzzaqueux. Each distribution was evaluated with Debye & Hückel correction for ionic strength. Neutralization by NaOH (virtual concentration “20 M” used in calculation, to make volume increase negligible) of 10 mM of compound in the presence of 160 mM of NaCl was chosen in order to approach the ionic strength value in electrophysiological experiments. The respective percentages at various pH values of the compound species H_2_A, HA^-^ and A^2-^ are indicated, except in the case of Asp where the H_3_A^+^, H_2_A and HA^-^ species are considered. Correspondence between the nomenclature used in this report and the reference nomenclature for carboxylic acids: malonic (propanedioic), succinic (butanedioic), glutaric (pentanedioic); fumaric ((*E*)-but-2-enedioic), maleic ((*Z*)- but-2-enedioic); *gamma*-aminobutyric acid (GABA) (4-aminobutanoic); aspartic (2-aminobutanedioic), (L)-glutamic ((*S*)-2-aminopentanedioic); malic (2-hydroxybutanedioic); oxaloacetic (OAA) (2-oxobutanedioic), α-ketoglutaric (2-oxopentanedioic); citric (3-hydroxy-3-carboxypentanedioic).

|  |  |  | pH<br>4.0 | pH<br>5.0 | pH<br>5.5 | pH<br>6.0 | pH<br>7.5 |
| --- | --- | --- | --- | --- | --- | --- | --- |
| Compound | Structure | pKa1;<br>pKa2 | H<sub>2</sub>A<br>HA<sup>-</sup><br>A<sup>2-</sup> | H<sub>2</sub>A<br>HA<sup>-</sup><br>A<sup>2-</sup> | H<sub>2</sub>A<br>HA<sup>-</sup><br>A<sup>2-</sup> | H<sub>2</sub>A<br>HA<sup>-</sup><br>A<sup>2-</sup> | H<sub>2</sub>A<br>HA<sup>-</sup><br>A<sup>2-</sup> |
| **Succinic acid**<br>SUCCIN |  | 4.3; 5.7 | 53<br>44<br>3 | 7<br>58<br>35 | 2<br>33<br>65 | 0<br>14<br>86 | 0<br>1<br>99 |
| **Fumaric acid**<br>FUMAR |  | 3.1; 4.5 | 3<br>40<br>57 | 0<br>6<br>94 | 0<br>2<br>98 | 0<br>1<br>99 | 0<br>0<br>100 |
| **1,2-(ditetrazol-5-yl)ethylene**<br>1 |  | 3.7; 5.1 | 18<br>21<br>61 | 1<br>22<br>77 | 0<br>8<br>92 | 0<br>3<br>92 | 0<br>0<br>100 |
| **N<sup>1</sup>,N<sup>4</sup>-dihydroxy fumaramide**<br>3 |  | > 8 | 100<br>0<br>0 | 100<br>0<br>0 | 100<br>0<br>0 | > 99<br>< 1<br>0 |  |
| **Dimethyl fumarate** |  | - | - | - | - | - | - |
| **Maleic acid**<br>MALE |  | 1.8; 6.6 | 0<br>98<br>2 | 0<br>90<br>10 | 0<br>74<br>26 | 0<br>47<br>53 | 0<br>3<br>97 |
| **Malonic acid**<br>MALON |  | 2.9; 5.7 | 5<br>91<br>5 | 1<br>66<br>34 | 0<br>38<br>62 | 0<br>16<br>84 | 0<br>1<br>99 |
| **Glutaric acid**<br>GLUTAR |  | 4.3; 5.4 | 59<br>37<br>4 | 7<br>46<br>47 | 2<br>23<br>75 | 0<br>9<br>91 | 0<br>0<br>100 |
| **Malic acid**<br>MAL |  | 3.5; 5.1 | 15<br>71<br>14 | 1<br>32<br>67 | 0<br>13<br>87 | 0<br>5<br>95 | 0<br>0<br>100 |
| **Aspartic acid**<br>Asp |  | 1.9; 3.7<br>pKa3=9.7 | 0<br>22<br>78 | 0<br>3<br>97 | 0<br>1<br>99 | 0<br>0<br>100 | 0<br>0<br>97 |

### Compound solutions

Stock di-CBX solutions (80 mM) were prepared at pH 5, with the aim to keep approximately constant the Na^+^ ion final concentration (165 + 10 + 6 = 181 mM), and 2 mL aliquots were kept at -20 °C for up to six months. Di-carboxylic acids were dissolved in water (about two third of the final stock volume) with partial neutralization to reach pH 5.0 using NaOH. Then enough 10x extracellular solution at pH 7.8 was added (some with and some without NaCl, in order to add (165 – 80*ρ) mM NaCl, depending on the counter-cation/di-CBX *ratio* ρ at pH 5), then pH and volume were re-adjusted using HCl and water. Before recording, a di-CBX aliquot (80 mM at pH 5) was diluted with 1x extracellular solution at the test pH and/or at pH 7.5, to reach final volume at the test pH. Then two #0.5 mL aliquots of the 1x di-CBX solution were isolated, supplemented with a drop of either 2 M HCl or 2 M NaOH, and used to finely adjust the di-CBX solution pH to the pH measured in the control (no compound) solution. Several pH measurements and adjustment were done (solutions without/with the di-CBX), rinsing the pH electrode with some control solution at the test pH, until the two pH values measured were equal within 0.01 pH unit. This was necessary because the test pHo 5.0 is near pHo50 (EC50), with maximal slope in the proton – GLIC activation curve (and di-CBXs have already no effect at pH 5.5). The di-CBX species distributions (given in Table 1) and counter-cation concentrations at pH 5.0 (and ρ) were evaluated (with Debye & Hückel activity coefficients consideration), for a solution of 10 mM di-CBX and 170 mM KCl, neutralized with “NaOH 20 M” (for calculation), using the free software Dozzzaqueux (Jean-Marie BIANSAN, http://jeanmarie.biansan.free.fr/dozzzaqueux.html, in French).

Picrotoxinin (Sigma P8390) [and caffeic acid] were dissolved in DMSO at 0.1 M [and 1 M] concentrations respectively, aliquoted (10 µL) in the bottom of 0.5 mL tubes, and kept at -20°C for up to a year. A 5-10 µL [or 1-2 µL] sample was diluted directly to 0.1 mM, by slowly adding it to the recording solution under fast bar stirring. An aliquot was sometimes centrifuged, re-frozen and re-used within three days. DMSO 0.1 % [or 0.01 %] had no effect on pHo 5-induced GLIC currents (whereas DMSO 1 % (0.14 M) produced a 15-20 % GLIC current inhibition). Solutions of the ditetrazole or dihydroxamic acid compounds were freshly prepared every day from the solid kept at -20°C. For each compound, two dissolution protocols were used, depending on days, before pH adjustment: the compound was either dissolved directly at 1 mM in the pH 7.5-recording solution through gentle bar stirring for 2-3 h at room temperature, or dissolved at 0.5 M in DMSO (within a minute) and then diluted under fast bar stirring, before pH adjustment. For each compound, the two dissolution protocols led to undistinguishable data.

### Experiments with oxaloacetate

The α-keto derivative of succinate, oxaloacetate (OAA), is known to be unstable in solution at low pH, due to spontaneous decarboxylation of its mono-anionic form (Tsai 1966). Indeed, after oxaloacetic acid was dissolved and partially neutralized with NaOH to reach pH 5.0 in our (Na^+^ compensated-) extracellular solution, as expected, we observed a progressive increase with time of the pH of this solution in the following minutes and hours. Therefore, particular care was taken to test OAA for an effect on GLIC at pHo 5.0: for each electrophysiological test, a solution of OAA was freshly prepared during whole-cell recording, and the solution at pH 5.0 was immediately loaded in the perfusion system, in order to allow a functional test within 3-4 minutes after dissolution. In this protocol, we observed that the OAA solution produced initially a potentiation of pHo 5-induced GLIC current. The increase in current vanished from a test to the next one, and turned into a decrease in GLIC current, actually due to a pH error increasing with time, the pH of the solution applied being more and more above the desired test-pHo value. The OAA data given in Fig. 2*EF* corresponds to the first test with OAA on each cell.

### Pharmacology: binding sites

The word “ortho*topic*” is used throughout this paper (instead of the more commonly used term “orthosteric”) to designate the neurotransmitter binding site of synaptic pLGICs, and the homologous site, situated “at the same *location*”, in GLIC or any pLGIC, as well as agents acting through this site. Similarly, “allo*topic*” (mentioned in Neubig *et al*. 2003) is used to comment a site situated at a *location* different from the reference, ortho*topic* site location. The word “allo*steric*” (Monod *et al*. 1965) is used to comment conformational transitions (*i*. *e*. 3D *steric* changes). A positive [or negative] allosteric modulator (PAM [or NAM]) modulates the allosteric transitions, towards [or away from] the active state (= open state). Fumarate in GLIC is a PAM acting by binding to the orthotopic site (an orthotopic PAM). Caffeate is an orthotopic NAM. In the associated report (Van Renterghem *et al*. Manuscript M2), we look for a contribution of allotopic binding (binding at the intra-SU pocket) to PAM or NAM effects. The proposed terminology has the advantage of not using the same word (“allosteric”) to designate two different concepts (both present in Monod, Wyman and Changeux, 1965): *other* binding site location and conformational transitions between active/inactive states.

1A/ Orthotopic site. The region in GLIC homologous to the pLGIC neurotransmitter binding site includes: Arg105 (Loop E), Arg133 (Loop B), Asp177 (Loop C), Asp181 (Loop C), and other residues, and is called in this report the “<u>orthotopic site</u> in GLIC”. 1B/ The <u>inter-subunit</u> (carboxylate) <u>binding pocket</u> in GLIC (Sauguet *et al*. 2013, Fourati *et al*. 2015, 2020) includes: Arg77 (Loop A), Arg105 (Loop E), Asp181 (Loop C), and, for the di-carboxylates, Asn152 (Loop F). The inter-SU pocket is therefore overlapping (Arg105, E181) with the conserved orthotopic site, but inter-SU compounds in GLIC are located slightly more deeply (than agonists in Eukaryote pLGICs). GLIC inter-SU pocket is accessible from the periphery of the pentamer through an entrance which is part of the orthotopic site (Arg133, Asp177). 2/ The <u>intra-subunit</u> (carboxylate binding) <u>pocket</u> in GLIC (Nury *et al*. 2010, Sauguet *et al*. 2013, Fourati *et al*. 2015, 2020) includes: Arg77 in its apo-GLIC conformation, and Arg85 (Pre-β5), Tyr102 (β6), Glu104 (β6). The intra-SU pocket, accessible from the vestibule *lumen*, corresponds exactly to the vestibular pocket in ELIC, and constitutes an “allotopic” binding site. When discussing (1/) *vs* (2/), we occasionally use “orthotopic” to designate the ensemble (1B+1A) (inter-subunit CBX pocket + its peripheral entrance), as opposed to the “allotopic”, intra-subunit (= vestibular) pocket (2/).

The word “impact” is used for changes in the receptor-channel protein (impact of a mutation, etc.), to distinguish from pH or compounds’ pharmacological “effects”. Unless otherwise stated, the notations FUMAR, (SUCCIN, etc.) and CBX, and the words succinate (caffeate, etc.) and carboxylate are used for the mixture of several acido-basic conjugated species in solution (see Table 1).

### Pharmacology: protocols and measurements

In most cases, a *protocol with pre-stimulation* was used: compound application started after GLIC stimulation at the same low pHo value (pHo 5, or another low-pHo value in Figs. 4, 5*CDF*). Pre-stimulation duration was 20 s (Figs 1*AB*, 2*EF*, 3*AB*, 4 *AB*, and two third of data in Figs 7, 8), except for: 10 s in the series of experiments with ditetrazole (Fig. 5*C*-*F*), dihydroxamic and other fumarate analogs; and 60 s in early experiments including compound selectivity evaluation (Fig. 2*A*-*D*), and about one third of the cells tested for residue dependency analysis (Figs. 7, 8). In Figs.7 and 8, 20 s- and 60 s-pre-stimulation data (in otherwise equal conditions) were pooled for analysis. Compound application had the same duration as low-pHo pre-stimulation. A *direct protocol* was used in some cases (Figs. 1*CD*, 3*C-F*): the compound solution at low-pHo was applied directly after the standard solution at pH 7.5.

**Figure 1.**
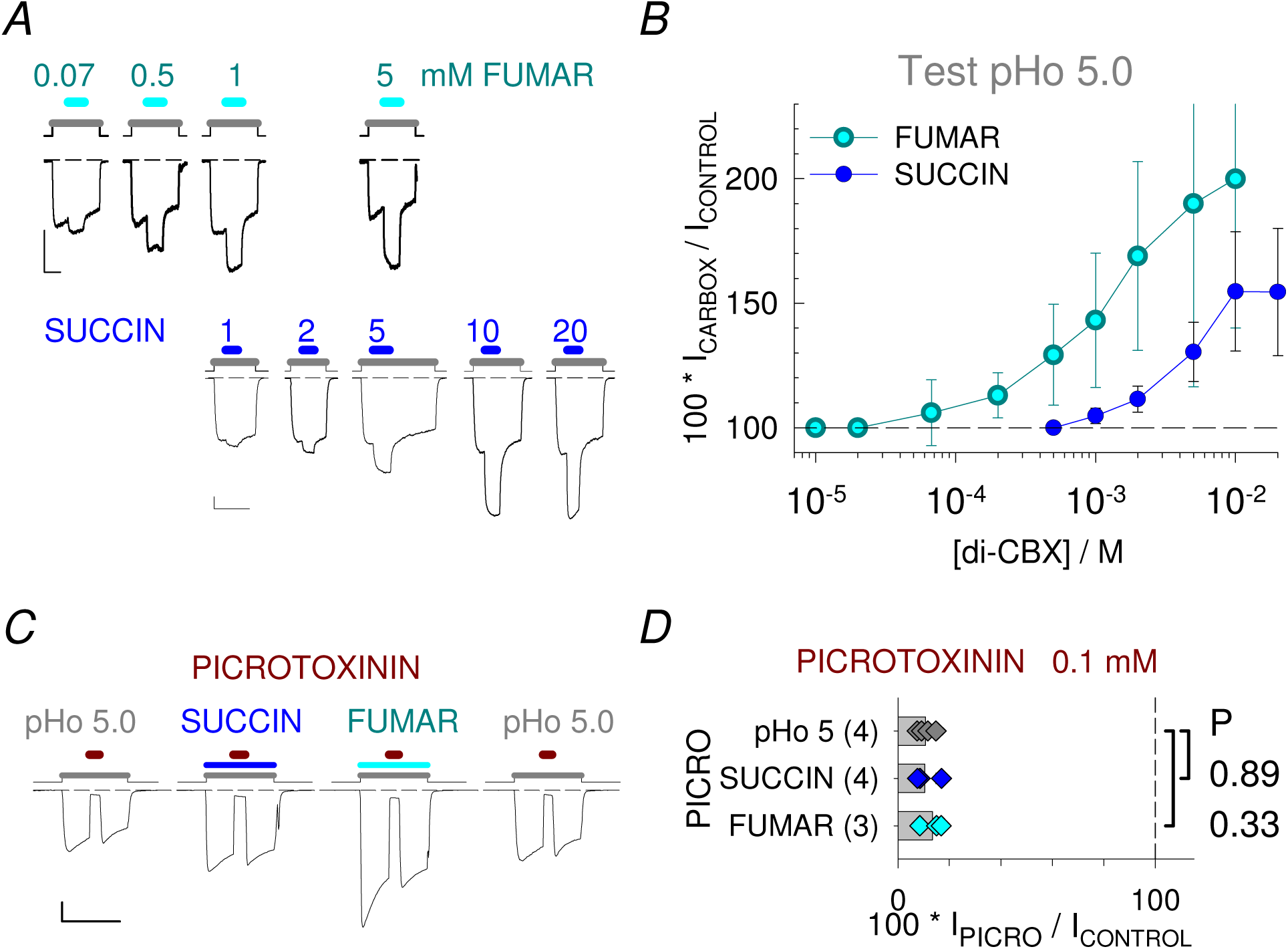
Fumarate and succinate are positive modulators of allosteric transitions in GLIC. *A*, *B*, ***Concentration-dependent potentiation by FUMAR or SUCCIN***. *A*, Whole-cell voltage-clamp current traces from tk-ts13 cells driven to express GLIC, showing the inward current elicited when pHo is changed from 7.5 to 5.0 (*Grey line*), and concentration-dependent potentiation by FUMAR (upper traces, *Cyan line*) or SUCCIN (lower traces, *Blue line*) applied as indicated (mM). *B*, Plots of concentration to effect, obtained with FUMAR and SUCCIN, in percent of the control current measured immediately before compound application. *C*, *D*, ***Inhibition of the di-CBX-elicited current by picrotoxinin***. *C*, Current traces showing the effect of picrotoxinin (0.1 mM; *Dark red line*) applied during stimulations at pHo 5, or during direct applications of SUCCIN or FUMAR, 10 mM at pHo 5. *D*, Corresponding bar graph of current in the presence of picrotoxinin, in percent of current before picrotoxinin application. Scale bars: 0.2 nA, 60 s.

Active current (I) was measured in reference to the current recorded at pHo 7.5 in patch-clamp, or 8.0 in oocyte experiments, and had negative values at the holding potential used (-40, -30 or -20 mV). Potentiation/inhibition was evaluated as [current in the presence of the compound] in percent of the [control GLIC–current] value. In the case of picrotoxinin (Fig. 1*CD*), the [control GLIC– current] was the [FUMAR@pHo5–stimulated current] measured immediately before toxin application. When the protocol with pre-stimulation was used (Figs 1*AB*, 2, 3*AB,* 4*AB,* 5*CF,* 7, 8), the [control GLIC–current] was the [low-pHo– stimulated current] measured immediately before compound application (except for some inhibition cases with correction for current decay, see below). Value of the [current in the presence of a compound] was measured at the early (negative) peak if any positive modulation was detected, and at the current minimum absolute value if not (picrotoxinin and caffeate; inactive compounds on WT GLIC (Fig. 2*D, F*); di-CBXs on most CBX-pockets mutants (Figs. 7, 8); compounds tested at pHo 6 (Figs. 4, 5*CD*)). When the inhibited-to-control % value measured in this way was comprised between 70 and 100 %, a correction was applied to the [control GLIC–current] value, in order to take into account GLIC current decay in the estimation of the control value. This estimation was done using one (or up to three) long-duration control (low pHo) application(s) on the same cell, and/or using an interpolation between current before compound, and current after compound wash-out. A similar correction of the control value was applied in the case of inhibitors in some cells where GLIC current decay was estimated to be “too fast”. No such correction was applied in measurements of potentiated-to-control % values, since potentiation was always relatively “fast”. When the direct protocol was used for FUMAR with various pipette solution pH values (Fig. 3), the current value measured was the peak value of either [current evoked by FUMAR at pHo 5] (Ipk[FUMAR@5]), or [current evoked at pHo 5] (Ipk[pHo 5]). Mutant and WT GLIC sensitivity to extracellular “protons” (H^+^ or H_3_O^+^ ions) was evaluated by measuring the peak value of the active current elicited by switching to low pHo (Ipk[pHo]). The data plot, Ipk[pHo] *vs* a_H+_o (or pHo), was fitted according to the Hill model at equilibrium (n.L+P = PL_n_, a all-or-none binding transition [of n ligand L on a protein P], with [P_total_] = [P]+[PLn], and with the dissociation constant arbitrarily noted K^n^). The equilibrium condition, written K^n^ = [L]^n^.[P]/[PL_n_], gives the Hill equation [PL_n_]/[P_total_] = 1/(1+K^n^/L^n^), here 1/(1+(a_H+50_)^n^/(a_H+_)^n^), also written [PL_n_]/[P_total_] = 1/(1+10^[n*(pH-pH50)]^) when expressed as a function of pH. Fit of “proton dose-response curve” electrophysiology data was done assuming I_pk_/I_pk_max = [PL_n_]/[P_total_], using the equation I_pk_ = I_pk_max/(1+ (a_H+50_)^n^/(a_H+_)^n^) for the patch-clamp data plotted as peak current *vs* proton activity, or the equation I_pk_ = I_pk_max/(1+10^[n*(pHo-pHo50)]^) for oocyte data plotted as peak current *vs* pHo. The data from each cell was fitted to give one pHo50 value. The parameters given in Fig. 6*C* (oocyte data) are the pHo_50_, [pHo_50_ standing for EpHo_50_, the pHo eliciting a half of the maximal activation reached using low pHo], and a ΔpHo_50_ from WT value, for which the pHo_50_ from each individual cell expressing a mutant was compared to the (mean) pHo_50_ from one (or two) cell(s) expressing wt GLIC recorded on the same day. Values are given as mean ± standard deviation, the value for each mutant is established from 3-6 cells coming from at least two injections. We occasionally use the notation EC_50_ (instead of an unusual “Ea_H+_o_50_”) to designate the extracellular solution proton activity eliciting half-maximal activation [despite the fact that we deal with a proton activity (*i*.*e*. 10^-pHo^, known from a pH-meter), whereas EC_50_ deals with a ligand concentration value in mol.L^-1^ (known from compound mass and solution volume)].

### Statistics

All values are given as mean ± standard deviation, of values obtained from individual cells. In the mutational analysis, the sample of values for each mutant was compared to the sample for WT GLIC using a Student’s T-test. The probability (P) for mutant and WT samples coming from a single normally distributed population is indicated near the mutant data, and P < 0.05 was the criteria used in text for a significant difference between samples.

### Chemistry

Briefly, the tetrazole bioisostere **1** was synthesized in 81% yield according to the method described by Chafin *et al*. (2008). The hydroxamic bioisostere **3** was obtained in a two-step procedure in an excellent 74% overall yield: *tert*-butyl O-protected hydroxylamine was coupled to fumaric acid in the presence of 1-(3-dimethylaminopropyl)-3-ethylcarbodiimide (EDCI) in water to give the *tert*-butyl O-protected intermediate **2** (Kapa *et al*. 2004). The protecting group was then removed under standard conditions to yield the desired hydroxamic compound **3**.

Commercially available reagents were used throughout without further purification. Analytical thin layer chromatography was performed on Merck 60F-254 precoated silica (0.2 mm) on glass and was revealed by UV light. ^1^H and ^13^C NMR spectra were recorded at 300 MHz and 75 MHz, or at 400 MHz and 100 MHz. The chemical shifts for ^1^H NMR were given in ppm downstream from tetramethylsilane (TMS) with the solvent resonance as the internal standard. Infrared (IR) spectra were obtained as neat films. High-resolution electrospray ionization mass spectrometry (ESI-HRMS) analysis was performed with a time-of-flight mass spectrometer yielding ion mass/charge (*m*/*z*) *ratios* in atomic mass units. For aromatic derivatives, purity was determined by reverse phase HPLC using a 150 mm x 2.1 mm (3.5 µm) C18-column: the compound was either eluted (method A) over 20 min with a gradient from 95% ACN/ 5% (H_2_O+0.1% HCO_2_H) to 100% (H_2_O+0.1% HCO_2_H) or (method B) over 5 min with 99% ACN/ 1% (H_2_O+0.1% HCO_2_H) then over 15 min with a gradient from 99% ACN/ 1% (H_2_O+0.1% HCO_2_H) to 100% (H_2_O+0.1% HCO_2_H).

<u>Preparation of (*E*)-1,2-di(2*H*-tetrazol-5-yl)ethylene (**1**)</u>: to a solution of (2*E*)-butenedinitrile (496 mg, 6.4 mmol, 1 eq.) in water (5 mL) were added sodium azide (922 mg, 14.2 mmol, 2.2 eq.) and zinc bromide (2.85 g, 12.7 mmol, 2 eq.). The reaction mixture was refluxed overnight. After cooling, concentrated HCl (1 mL) was added at 0°C and the reactive was stirred for 2 h then filtered and the residue washed with water to give the desired compound **1** as a raw solid (848 mg, 81% yield); mp = 276.5 °C (deflagration), in accordance with published values (Demko *&* Sharpless 2001); ^1^H NMR (300 MHz, DMSO-*d6*) δ 7.67 (s, 2H, CH=CH), 12.80 (bs, 2H, 2 NH tetrazole); ^13^C NMR (75 MHz, DMSO *d6*) δ 119.5 (CH=CH), 153.6 (Cq tetrazole); IR (cm^-1^) ν 3095, 3041, 1547, 1082, 1068, 1010, 858; HRMS (TOF MS ES+) *m/z* calcd for C_4_H_5_N_8_^+^ (M+H)^+^165.0632 found 165.0640; HPLC purity 100% t_r_ = 11.50 min (Xbridge C18 column, 254 nm, method B).

<u>Preparation of di-*tert*-butyl fumaroyldicarbamate (**2**)</u>: to a solution of fumaric acid (235 mg, 2.0 mmol, 1 eq.) in water (20 mL) were added O-*tert*-butylhydroxylamine hydrochloride (553 mg, 4.4 mmol, 2.2 eq.), sodium hydroxide (177 mg, 4.4 mmol, 2.2 eq.) and, after 30 min, 1-(3-dimethylaminopropyl)-3-ethylcarbodiimide hydrochloride (958 mg, 5.0 mmol, 2.5 eq.). After stirring at room temperature for 16 h, the reaction mixture was filtered and the solid was recrystallized in methanol to yield the desired compound **2** as a white solid (452 mg, 87% yield). mp = 275 °C (decomposition); ^1^H NMR (300 MHz, CD_3_OD) δ 1.28 (s, 18H, 2 OC(CH_3_)_3_), 6.93 (s, 2H, CH=CH); ^13^C NMR (75 MHz, CD_3_OD) δ 26.6 (OC(*C*H_3_)_3_), 83.9.0 (O*C*(CH_3_)_3_), 131.6 (CH=CH), 164.9 (CO); IR (cm^-1^) ν max 3236, 2979, 1671, 1057; HRMS (TOF MS ES+) *m/z* calcd for C_12_H_23_N_2_O_4_^+^ (M+H)^+^ 259.1652 found 259.1656.

Preparation of *N*^1^,*N*^4^-dihydroxyfumaramide (**3**): a solution of compound **2** (300 mg, 1.2 mmol) in a TFA/DCM 1/1 mixture (20 mL) was refluxed overnight. The reaction mixture was filtered to yield the desired compound **3** as a white solid (145 mg, 85% yield). mp = 231 °C (decomposition); ^1^H NMR (400 MHz, DMSO-*d6*) δ 6.70 (s, 2H, CH=CH), 8.00-10.00 (bs, 2H, 2 OH), 10.00-11.50 (bs, 2H, 2 CONH); ^13^C NMR (100 MHz, DMSO *d6*) δ 129.5 (CH=CH), 160.8 (CO); IR (cm^-1^) ν max 3C_4_H_7_N_2_O_4_^+^ (M+H)^+^ 147.0400 found 147.0401; HPLC purity 82% t_r_ = 1.84 min (Xselect C18 column, 254 nm, method A).

### Determination of acidity constants

For (*E*)-1,2-di(2*H*-tetrazol-5-yl)ethylene (Compound **1**): A solution of 15.9 mg (around 0.1 mmol) in DMSO (1 mL) was diluted with deionized water to reach 100 mL of stock solution. 20 mL were then titrated by a 0.02 M NaOH solution at 25°C under pH measurement. The curve pH=f(V_NaOH_) allowed a precise determination of Veq_2_ (volume of NaOH used to neutralize the two acidities of Compound **1**). Values p*K*a_1_ = 3.7 and p*K*a_2_ = 5.1 were determined from the curve as equal to pH values at (0.25 x Veq_2_) and (0.75 x Veq_2_) respectively. For Compound **3**, p*K*a values could not be determined by this method because of the known poor acidity of hydroxamic acids. They were estimated above 8 considering known values for hydroxamic acids (Singh 2015).

## Results

### 1/ Fumarate is a positive modulator of GLIC

Using whole-cell recording, and a *protocol with pre-stimulation*, in which the GLIC-expressing cell was first exposed to an extracellular solution at pH 5, we show that FUMAR potentiates pHo 5-activated GLIC current (Fig. 1*A*, *upper trace*). The pHo 5 value is known to produce approximately a half of the maximal activation reached using low-pHo (Bocquet *et al*. 2007, Nury *et al*. 2011, Nemecz *et al*. 2017). The 4-carbon di-CBX applied (10 mM) at the control, sub-threshold pHo values 7.5, 7.0, 6.5 had no effect, showing that FUMAR has no agonist effect. Potentiation by FUMAR was reversible and concentration dependent, with a threshold concentration near 0.1 mM at pHo 5, and a maximal effect near 10 mM (Fig.1*B*). SUCCIN also produced a reversible and concentration dependent potentiation of GLIC current (and no agonist effect), but with approximately a one order of magnitude right shift of the concentration-effect curve, and a reduced maximal PAM effect (Figs. 1*A lower trace* and 1*B*).

In order to ascertain that the increase in inward current induced by FUMAR or SUCCIN originates in GLIC modulation, we checked some pharmacological and ion selectivity properties. For this we used a *direct protocol*, where the control extracellular solution at pH 7.5 was replaced directly by a solution containing FUMAR or SUCCIN at pHo 5 (FUMAR@5, SUCCIN@5). Applied at 10 mM on mock-transfected tk-ts13 cells, FUMAR@5 (n=2), as well as SUCCIN@5 (n=2), did not produce more current than pHo 5 alone (-6.4 (± 2.8, n=6) pA at - 50 mV). The contribution of endogenous channels/transporters to pHo 5-elicited currents was not taken into account in our studies on GLIC at negative membrane voltage. On GLIC-expressing cells, the SUCCIN@5-and FUMAR@5-activated currents were almost fully inhibited by picrotoxinin (0.1 mM), as was pHo 5-activated current (Fig. 1*CD*). These properties show that SUCCIN@5- and FUMAR@5-induced currents depend on current flow through GLIC pentamers in the membrane.

The reversal potential (Erev) for [FUMAR@5]-induced current exhibited the behavior that we commonly observe when a large GLIC current is recorded from a whole cell. In our recording conditions [using sodium outside (Na^+^ 181 mM, independent of pHo and FUMAR concentration, see Methods), and caesium inside (Cs^+^ #188 mM at pHi 7.3), and using a ramp protocol], GLIC current ([pHo 5] or [FUMAR@5]) reverses at a positive value >+10mV at first ramp. But Erev is drifting down from one ramp to the next one (up to 5 mV decrease within 2 s between first and second ramp), due to the accumulation of Na^+^ ions at the cytosolic end of GLIC ion pores, ending in quasi symmetrical Na^+^ conditions. This Erev shift-down is reversible within a few minutes of washout. On three cells tested, the initial Erev values observed were +13.0 (± 1.1, n=3) mV with [FUMAR@5] (10 mM), and +12.5(±2.2, n=3) with pHo 5 three minutes later. Assuming equal activity coefficients for Na^+^ outside and Cs^+^ inside, these underestimated Erev values both correspond to a permeability *ratio* P_Cs_/P_Na_ of 0.58. From other experiments with low-pHo activation (including single channel recording at various constant membrane voltage values), we observed that the actual Erev is near +15 mV (P_Cs_/P_Na_ near 0.54). But from our observations with the ramp protocol, we conclude that the [FUMAR@5] elicited current has the signature of a current flowing through GLIC ion pores, which are permeable to Cs^+^ ions, but to a lesser extent than to Na^+^ ions.

From this ensemble of data, we conclude that SUCCIN and FUMAR, applied extracellularly, fill the criteria for a PAM of GLIC. The di-CBX compounds however do not activate GLIC at neutral pHo.

### 2/ Fumarate is the best positive modulator identified

**Figure 2.**
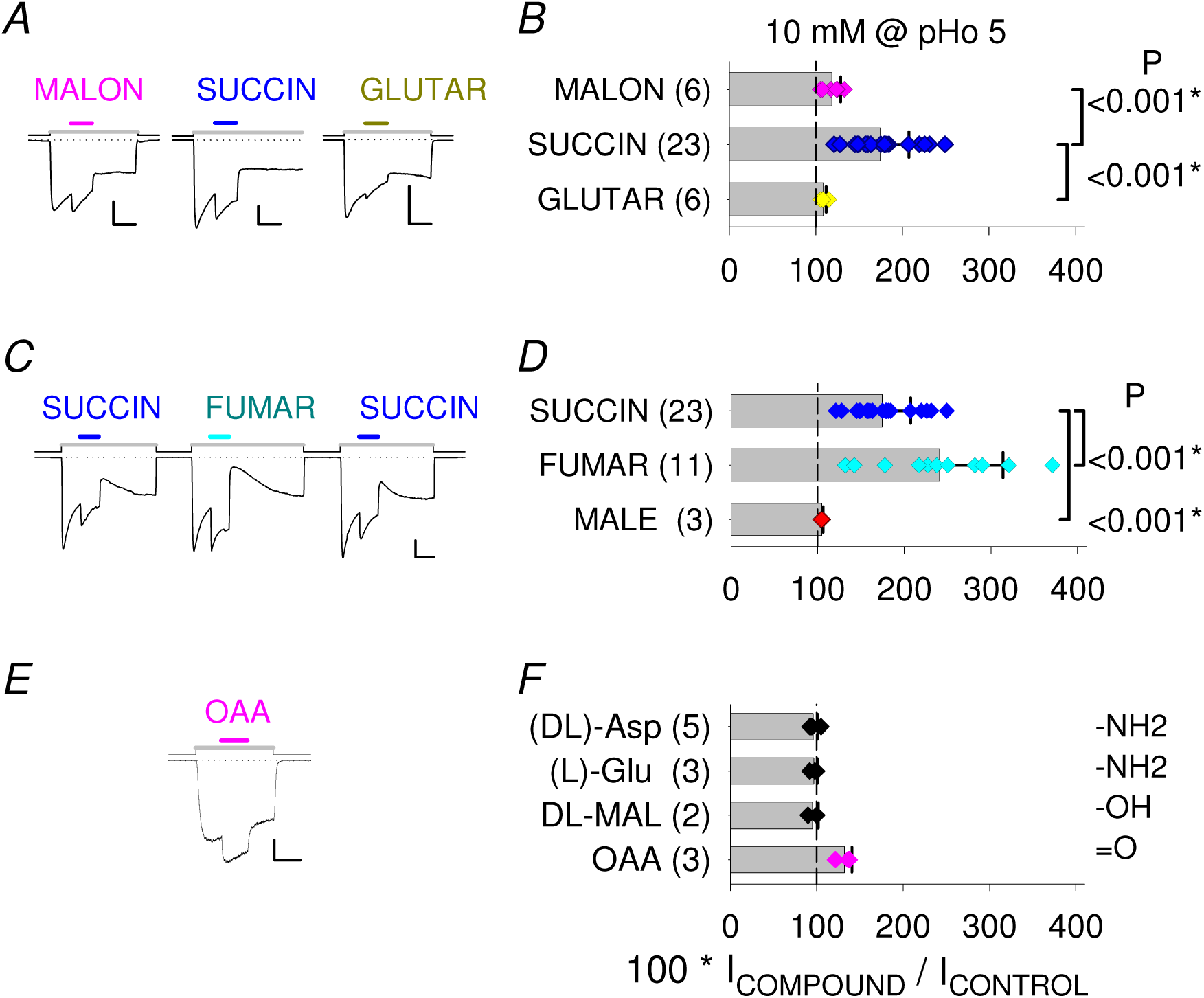
Fumarate is the best PAM identified. *A*, *B*, ***Optimal 4-carbon chain length***. *A*, Current traces from three cells, showing potentiation of the current elicited at pHo 5.0 (*Grey line*) by di-CBX compounds with three (malonate, *Left*, *Pink line*), four (succinate, *Middle*, *Blue line*) or five carbons (glutarate, *Right*, *Dark yellow line*), and *B*, corresponding bar graph. *C*, *D*, ***Optimal elongated shape of the 4-carbon di-CBX molecule***. *C*, Current trace showing that FUMAR (*trans* configuration; *Middle*) produces a larger potentiation than the saturated 4-carbon di-CBX SUCCIN (*Left* and *Right*) applied at the same concentration on the same cell. Note strong OUT effect in this recording. *D*, Corresponding bar graph. Any PAM effect is lost with the *cis* configuration of maleate (MALE; 10 mM). *E*, *F*, **Effect of neighbour compounds**. *E*, Current trace showing the potentiating effect of oxaloacetate (OAA; unstable in solution, see *Methods*; *Pink line*). *F*, Bar graph showing the PAM effect of the 2-oxo-derivative OAA, and absence of effect of the 2-amino- and 2-hydroxy-derivatives aspartate, glutamate and malate. All compounds (10 mM) were tested at pHo 5, in sets of cells distinct to those used for Figure 1 data. Scale bars: 0.5 nA, 60 s (*A*, *C*); 1 nA, 20 s (*E*).

Other compounds with two carboxyl groups were tested at 10 mM with pre-stimulation at pHo 5. Among saturated di-CBX compounds, malonate (MALON) and glutarate (GLUTAR) were active as PAMs but weaker than SUCCIN (Fig. 2*AB*), giving in this condition the order: 4-carbon > 3-carbon > 5-carbon. Among 4-carbon di-CBXs (Fig. 2*CD*), the rigid molecule FUMAR was more active than the flexible saturated compound SUCCIN, and the diastereomer maleate (MALE) was inactive (Fig. 2*CD*), showing that the *trans* double bond present in FUMAR optimizes the PAM action, whereas the *cis* double bond in MALE impairs any PAM efficiency. Among di-CBX derivatives, aspartate and glutamate (α-amino-succinate, α-amino-glutarate) were inactive, as well as malate (α-hydroxy-succinate) (Fig. 2*F*). Oxaloacetate (OAA; α-keto-succinate) was found to be active but unstable (Fig. 2*EF*; see also Methods) and was not further studied.

Additional compounds were also tested with pre-stimulation at pHo 5. The dipeptides Gly-Glu and *N*-acetyl aspartyl glutamate (1 mM) had no effect. The tricarboxylate citrate was tested at two concentrations (1 and 5 mM) at both pHo 5.0 and pHo 5.5, and showed no effect on GLIC current amplitudes, but produced some instability in the recording, manifested as small superimposed brief transient inward currents.

The decreasing order observed (FUMAR > SUCCIN > MALON > GLUTAR > MALE) shows that a 4-carbon chain length with the carboxyl groups in a *trans* configuration optimizes the PAM efficiency. FUMAR is the best PAM of GLIC identified in this study.

### 3/ GLIC activity and FUMAR potentiation *ratio* vary with intracellular pH

We commonly observe in various ways that the GLIC current, recorded with a given low pHo value, decreases when pHi decreases. In other words, the GLIC pentamer, well known to be activated at low pHo, is inhibited at low pHi. This property is already noted by Hilf and Dutzler (2009), in a Supplementary Figure reporting data apparently coming from a single inside-out macro-patch experiment, using solutions which are not described except for their pH. Although mostly forgotten in later GLIC literature, this data denotes an inhibition of GLIC current when pHi decreases, when using a stationary extracellular stimulation at pHo 4.

**Figure 3.**
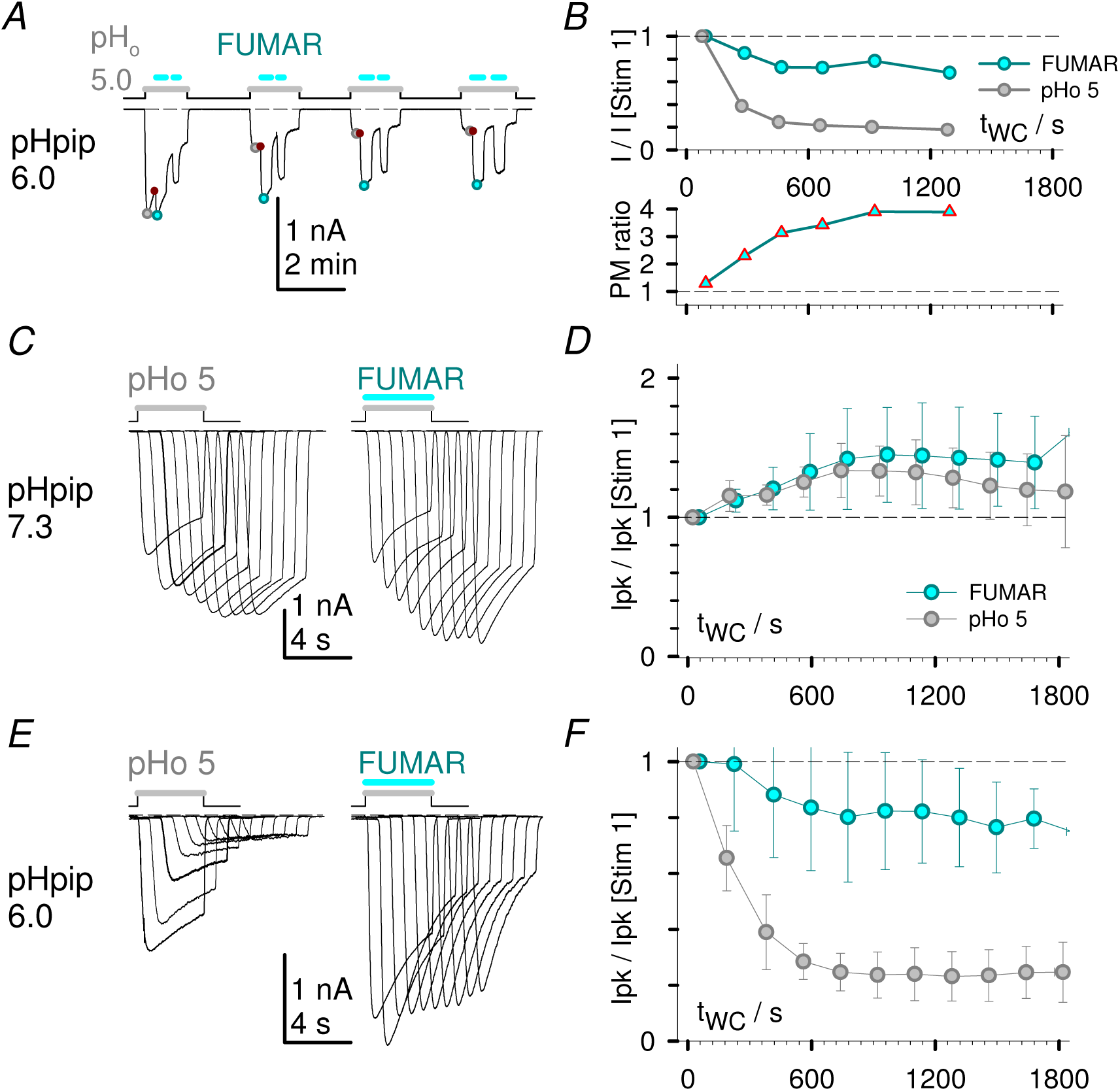
FUMAR-potentiated current (at pHo 5) is less reduced with a low-pH pipette solution than pHo 5-elicited GLIC current. *A*, *B*, ***Influence of a pH 6 pipette solution in a FUMAR experiment using a protocole with pre-stimulation.*** *A*, Continuous current trace showing the first twelve minutes of an experiment with two FUMAR applications/wash-out per stimulation at pHo 5. Wash time (pHo 7.5) was 92 – 112 s between stimulations. *B*, *Upper Graph*, decrease with [time spent in the whole-cell configuration] (t_WC_) of Ipk[pHo 5] and peak FUMAR current (measured at the first one of two FUMAR applications), both normalized to their respective values at first stimulation, and *B*, *Lower Graph*, increase with t_WC_ of the PAM ratio (here FUMAR peak current value, cyan circle in *A*, over pHo 5 current value measured immediately before FUMAR application, red triangle in *A*). *C*-*F*, ***Influence of the pipette solution pH in FUMAR experiments using a direct protocole.*** The pipette solution was either at pH 7.3 (*C*, *D*) or at pH 6.0 (*E*, *F*). *C*, *E*, Current traces recorded with repetitive pairs of 4 s-duration extracellular stimulations first with pHo 5 (*Grey line*; *Left* traces), then with [FUMAR 10 mM at pHo 5] (direct protocol; *Grey and Cyan lines*; *Right* traces). The 4 s-stimulations in a pair were applied every 30 s, and pairs were applied every 180 s. Traces corresponding to ten consecutive pairs are displayed superimposed, with a +0.7 s abscissa drift between pairs. *D*, *F*, Corresponding plots of t_WC_ *versus* peak current values Ipk[pHo 5] (*Grey circles*) and Ipk[FUMAR@5] (*Cyan circles*), normalized to their respective Ipk values in the first pair for each cell. Mean values from 3 cells with pHpip 7.3 (*D*) and 6 cells with pHpip 6.0 (*F*).

We report here data obtained using whole-cell recording with a pipette solution pH (pHpip) adjusted at either 7.3 or 6.0, in order to slowly modify the pHi value, while taking advantage of some delay in cell-pH relaxation to the pHpip value, following diffusion to and from the cytosol through the recording pipette orifice (Fig. 3). Using pHpip 6.0 and a protocol with pre-stimulation, the peak FUMAR-potentiated current recorded was almost constant from a test to the next one, whereas Ipk[pHo5] decreased, leading to a potentiation *ratio* increasing with time of whole-cell recording (see Fig. 3*AB*). We chose a direct protocol to further quantify the influence of pHpip, in order to reduce the tests duration (*i*. *e*. the proportion of time spent at low-pHo). Pairs of 4 s-extracellular stimulations, first with pHo 5, then with FUMAR at pHo 5 (direct protocol; start-to-start delay: 30 s), were applied every three minutes, in order to follow the variation of current amplitudes with time, *i*.*e*. with whole-cell dialysis duration. With the standard pipette solution (pHpip 7.3), the pHo 5-elicited current peak amplitude (Ipk[pHo 5]) became approximately stable with time (<30 min), after slightly increasing in the first ten minutes (Fig. 3*CLeft traces,* and 3*D*). But with a pipette solution adjusted at pH 6, Ipk[pHo 5] strongly decreased in the first ten minutes, before reaching a plateau at about one fifth of its initial value (Fig. 3*ELeft traces* and 3*F*). This data is consistent with the interpretation that a progressive relaxation of the cell-pH towards the pHpip value 6 (following diffusion of low-pH buffered solution from the pipette) causes a progressive decrease in pHo-elicited GLIC currents. Whereas cell-solution equilibration toward the control pipette solution (with pHpip 7.3 thought to be near a mammalian cell spontaneous pHi) is accompanied by some usual rise in Ipk[pHo 5] in the first 10-15 minutes, due to recording configuration improvement, and stabilization of permeant/non permeant ions concentrations.

Meanwhile, considering the second stimulation of each pair, the peak current elicited by FUMAR (Ipk[FUMAR@5]) showed the same kinetic variation as Ipk[pHo 5] when pHpip 7.3 was used (Fig. 3*CRight traces*, and 3*D*), but only a minor reduction to about four-fifths of its initial value when using pHpip 6 (Fig. 3*ERight traces* and 3*F*). This data shows that, when intracellular pH decreases, Ipk[pHo 5] decreases much more than Ipk[FUMAR@5], so that the FUMAR potentiation *ratio* strongly increases when pHi decreases.

This property is probably the main cause of the great variability observed in our FUMAR data at pHo 5 (see and compare Figs. 1*B*, 2*D*, 4*B*, 5*F*, from distinct cell samples): it may be that the sub-membrane pHi value was not fully controlled in the sets of experiments presented (as cell dialysis varies with numerous parameters, such as access conductance and cell volume, see Moroni *et al*. 2011). So that, most importantly, a variable drop in sub-membrane pHi, induced by the extracellular application of low-pHo stimulating solutions, may appear as the major determinant in the variability of both the FUMAR PAM effect, and the observed GLIC current decay kinetics (see Discussion).

### 4/ Influence of pHo on the FUMAR modulatory effect

**Figure 4.**
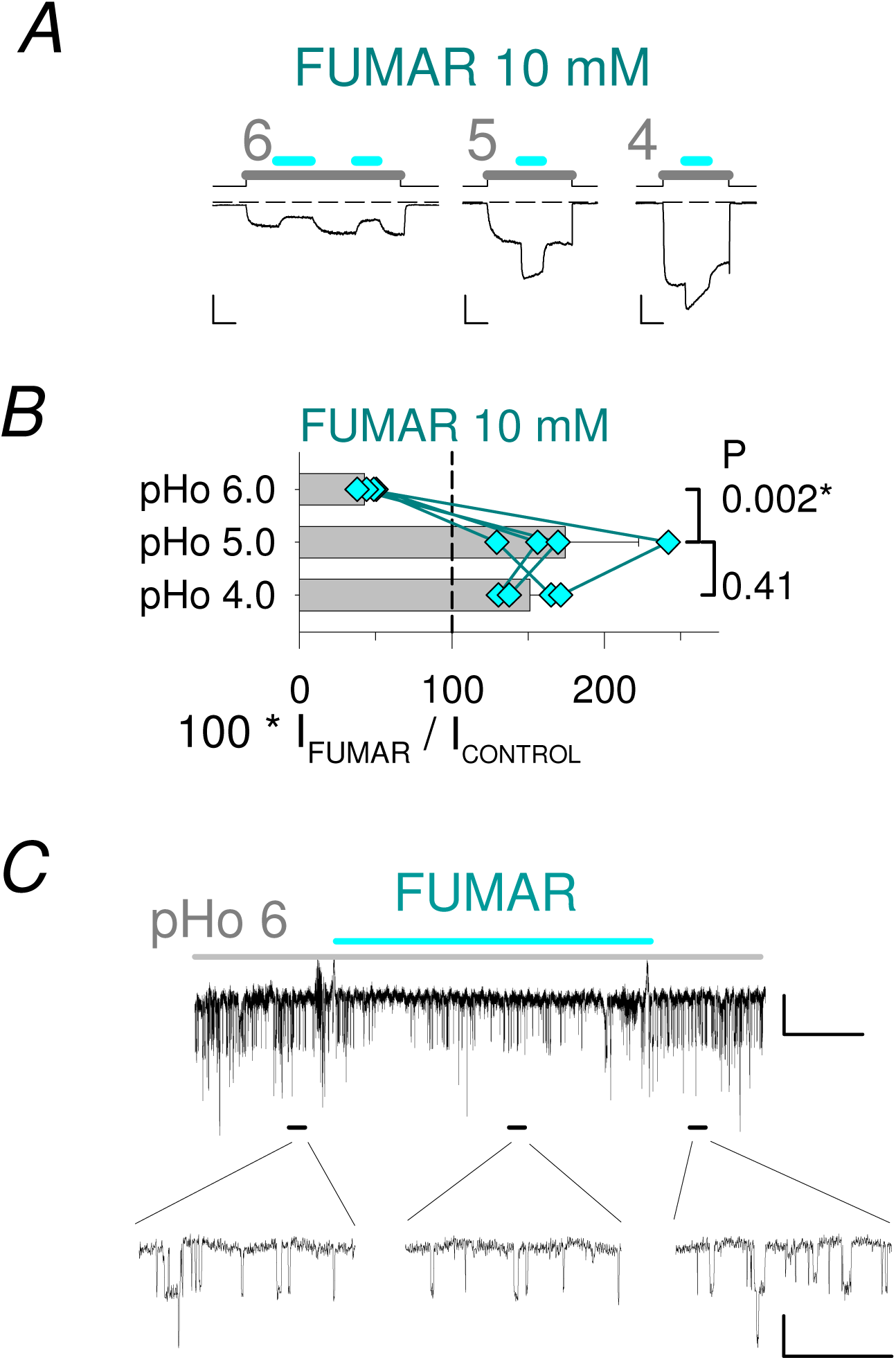
FUMAR PAM effect occurs only in the upper part of the proton activation curve. *A*, Whole-cell current traces showing opposite effects of FUMAR (10 mM; *Cyan line*) on the low-pHo elicited current, at pHo values 6.0 (*Left*, inhibition), or 5.0 and 4.0 (*Middle* and *Right*, positive modulation). Scale bars: 0.2 nA, 20 s. *B*, Corresponding bar graph. Data points obtained from the same cell (n=4) are joined by a line. *C*, Trace of single channel current recorded from an outside-out patch, showing the paradoxical negative modulation by FUMAR (10 mM) of GLIC activity at pHo 6.0, and time-expanded 2 s-duration details captured before, during and after FUMAR application, as indicated by black bars. Scale bars in *C*: 0.5 pA; 10 s (*Upper trace*) and 1 s.

Using our standard pHpip 7.3, surprisingly, the PAM effects of SUCCIN and FUMAR occurred only in a range of low pHo values (*i*. *e*. in the upper part of the GLIC activation curve). SUCCIN (10 mM), a positive modulator at pHo 5.0, had virtually no effect at pHo 5.5. FUMAR, more potent than SUCCIN at pHo 5, was chosen to further examine the influence of the test-pHo value. When FUMAR (10 mM) was applied after pre-stimulation at pHo 4.0, a potentiation of GLIC current was observed (Fig. 4*AB*). Potentiation was smaller at pHo 4.0 (151 % of control (± 20 %, *n*=4), than at pHo 5.0 (174 (± 48, *n*=4) % of control in this sample of cells), consistent with the expected decrease of a PAM effect when increasing the agonist concentration. Unexpectedly, the small pHo 6.0-activated inward current was reduced by FUMAR: the percentage of current in the presence of FUMAR (10 mM) was 43 (± 6, *n*=4, P = 0.002) % of control at pHo 6.0 (Fig. 4*AB*). Single-channel recording from an outside-out patch at pHo 6.0 allowed observing a decreased GLIC activity in the presence of FUMAR (10 mM; Fig. 4*C*). Therefore, FUMAR, a PAM at pHo 4-5, is unexpectedly a weak NAM at pHo 6.

### 5/ Synthesis and test of ditetrazole and dihydroxamic acid FUMAR bioisosteres

We were interested in using GLIC as a kind of bioassay to test the biocompatibility of structural analogs of the carboxyl group. We decided to build structural analogues of FUMAR, using a replacement of carboxyl groups by the bioisosteric tetrazolyl or hydroxamyl groups (Ballatore *et al*. 2013), and therefore synthesized bi-functional analogues of 4-carbon atoms length comprising an ethylene moiety substituted in *trans* configuration with either two tetrazole (compound **1**) or two hydroxamic acid functions (compound **3**) (see Fig. 5*AB* and Methods).

**Figure 5.**
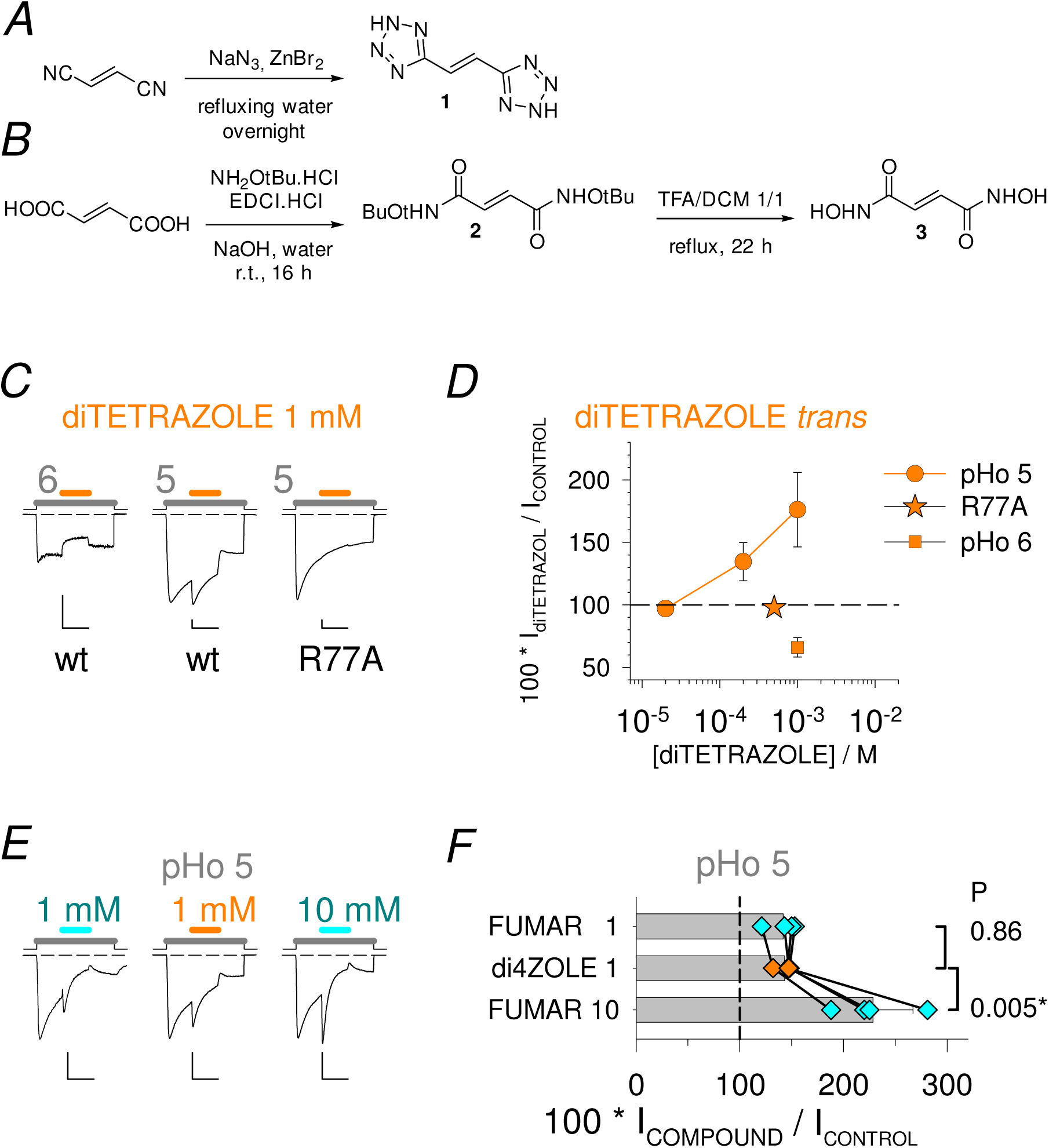
A ditetrazole FUMAR analog mimics FUMAR effect. *A*, *B*, ***Chemical synthesis of fumarate bioisosteres.*** Methods used to synthetize the ditetrazole compound **1** (*A*) and the dihydroxamic compound **3** (*B*) used in this study. See Methods for details. *C*, *D*, ***FUMAR-like effect of the ditetrazole compound 1***. *C*, Current traces showing inhibition at pHo 6 (*Left*) and potentiation at pHo 5 (*Middle*) of low-pHo elicited current, by the ditetrazole compound **1** (1 mM; *Orange line*), on cells expressing GLIC WT, and absence of potentiation (0.5 mM at pHo 5) on the R77A GLIC variant (*Right*). *D*, Plot of the ditetrazole compound **1** data with three concentrations at pHo 5 (n=6; *Circle*), and 10 mM at pHo 6 (n=2; *Square*), on GLIC WT, and with 5 mM at pHo 5 on GLIC R77A (n=2; *Star*). *E*, *F*, ***Comparison of ditetrazole and FUMAR effects on the same cell.*** *E*, Current trace from one cell, comparing the PAM effects at pHo 5 of the ditetrazole compound **1** (di4zole; 1 mM, *Middle*) and FUMAR (1 and 10 mM, *Left* and *Right*). *F*, corresponding data from 4 cells. Scale bars: 0.1 nA (*CLeft*) or 0.5 nA; 10 s.

The ditetrazole compound **1** (Fig. 5*A*) was active on GLIC, and displayed a PAM effect at pHo 5 (Fig. 5*C-F*). As with FUMAR, compound **1** PAM effect was suppressed following alanine substitution of GLIC Arg77 (see next section; Fig. 5*CD*). And the ditetrazole tested at pHo 6 reduced the pHo 6-elicited inward current (Fig. 5*CD*). This data suggests that the tetrazolyl group is efficiently recognized by GLIC carboxyl group binding sites, and that the ditetrazole **1** molecule is efficiently recognized by the FUMAR binding site(s) involved in the modulation of GLIC activity.

In contrast to the ditetrazole, the dihydroxamic acid FUMAR analogue **3** (Fig. 5*B*) (0.1 to 1 mM, from 2 dissolutions without DMSO and 2 dissolutions in DMSO) was inactive on pHo-elicited currents at both pHo 5 (n=8 cells) and pHo 6 (n=2) on GLIC WT, and up to pH 6.5 on the gain of function mutant GLIC T255A (n=1). We also tested on GLIC, at pHo 5 and pHo 6, a commercially available di-ester of FUMAR: dimethylfumarate (2 mM) was inactive on GLIC (n=3 each).

In this work with FUMAR structural analogues, we had some interest in comparing the pHo windows allowing a PAM effect, when using compounds with distinct species distributions with pH (in order to decide if the H_2_A compound species may be active, and the HA^-^ and A^2-^ species inactive, an hypothesis suggested by the FUMAR data, see Fig. 4). The results however did not allow this comparison. A measure of the ditetrazole **1** acidity constants was performed (see Methods), resulting in p*K*a1 and p*K*a2 values of 3.7 and 5.1, both between the respective p*K*a values of FUMAR and SUCCIN (see Table 1).

FUMAR, SUCCIN and the ditetrazole have therefore very similar species distributions with pH (Table 1). With p*K*a values expected to be larger than 8, the dihydroxamic acid isostere (compound **3**) is thought to be present only as a diprotonated species (H_2_A) at pHo 5.0 to 6.5, but it was inactive. Finally, dimethylfumarate, a putative analog of the FUMAR di-acid species (H_2_A) independent of pHo, was inactive as well. Therefore, our data with FUMAR analogs gives no information on whether the inversion of FUMAR effect on GLIC between pHo 5 and pHo 6 is due to a pH effect on the FUMAR molecule protonation state, or a pH effect on the GLIC ECD protonation state.

### 6/ Mutational analysis: orthotopic site/inter-SU site, and vestibular/intra- SU site in GLIC ECD

A mutational analysis was performed to establish the residue-dependency of the di-CBX PAM effects, using a series of single amino-acid GLIC variants. FUMAR and SUCCIN were both tested for comparison.

*Sites and single mutations chosen*. The putative sites evaluated (Fig. 6*AB*; see also the Methods section: *Pharmacology: Binding sites*) were chosen according to crystallographic data which characterized in GLIC ECD two acetate binding pockets (Sauguet *et al*. 2013, Fourati *et al*. 2015), also hosting other carboxylate compounds (Fourati *et al*. 2020). GLIC inter-SU CBX-binding pocket (Sauguet *et al*. 2013) overlaps with the region homologous to the conserved pLGICs’ orthotopic neurotransmitter binding site. But compounds in GLIC (*i*.*e*. the inter-SU pocket) are slightly more deeply buried (and slightly “lower” = closer to the membrane) than the Eukaryote orthotopic site ligands. In docking predictions in GLIC, both CAFFE (Prevost *et al*. 2013) and CROTON (Alqazzaz *et al*. 2016) were located slightly lower than the pLGICs reference site. And the inter-SU site slightly deeper location was confirmed in all available CBX-bound GLIC structures (Fourati *et al*. 2020). As a consequence, the inter-SU pocket in GLIC is accessible from the periphery of the pentamer through an empty peripheral entrance, which in Eukaryotes is part of the orthotopic site. 2/ The intra-SU CBX-binding pocket in GLIC ECD (Nury *et al*. 2010, Sauguet *et al*. 2013, Fourati *et al*. 2015, 2020) is accessible from the vestibule *lumen*, and corresponds exactly (see Fourati *et al*. 2015) to the *vestibular compound binding pocket* emphasized in ELIC (Spurny *et al*. 2012), and also occupied by a compound in sTeLIC (Hu *et al*. 2018). A cavity homologous to the prokaryotic vestibular compound binding pocket was identified in the crystal structures of several Eukaryote pLGICs, but it is then compound-empty, and either vacant or occupied by Loop Ω (Hu *et al*. 2018, Brams *et al*. 2020).

The amino-acids chosen were: for the vestibular, intra-SU pocket, (Arg77), Tyr102 (β6 strand) and Glu104 (β6 strand); for the inter-SU pocket, Arg77 (Loop A), Arg105 (Loop E), Asn152 (Loop F), and Glu181 (Loop C); and an additional two positions in the peripheral entrance to the inter-SU pocket: Arg133 (Loop B) and Glu177 (Loop C). We also included the GLIC position Tyr23, homolog to ELIC’s Phe19 used by Spurny *et al*. (2012) to conclude that the inhibitory effect of flurazepam on the agonist action of GABA is mediated by binding to the orthotopic site in ELIC.

**Figure 6.**
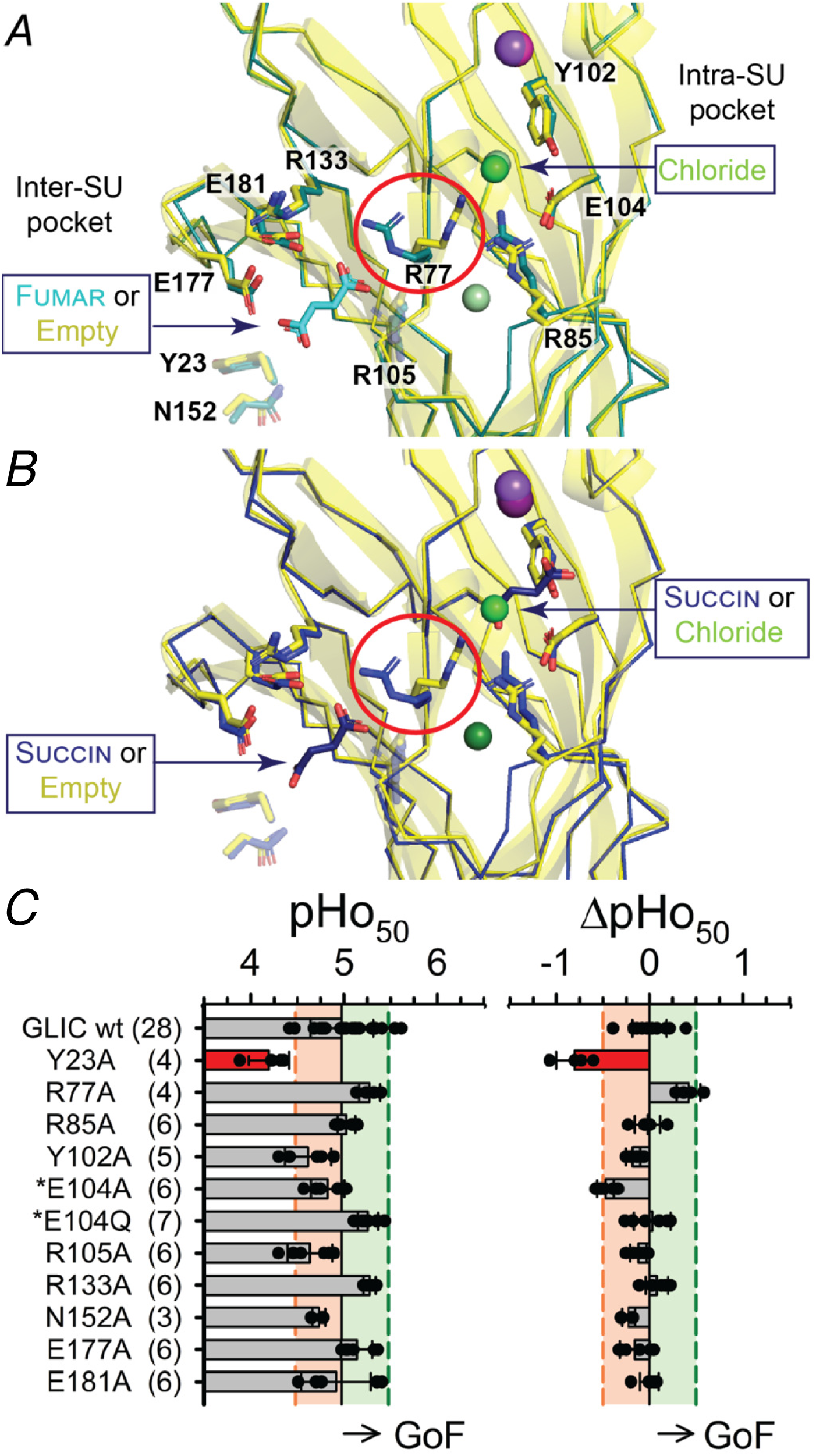
CBX-pockets single mutations: location in GLIC structure, and low impact on GLIC proton sensitivity. *A*, *B*, Structural position in GLIC ECD of the amino-acids considered in the mutational analysis, and inter-SU CBX-associated pivot movement of Arg77 side-chain. *Left*, inter-SU CBX-binding pocket (Arg77, Arg105, Asn152, Glu181), and its peripheral entrance (Arg133, Glu177). *Right*, intra-SU CBX-binding pocket (Arg77(apo), Arg85, Tyr102, Glu104). *A*, Superimposed views from the GLIC-FUMAR co-crystal structure (PDB reference 6hj3; *Dark* c*yan*) and the Apo-GLIC (GLIC in Phosphate buffer) structure (4qh5; *Yellow*), and *B*, superimposed views from GLIC-SUCCIN (6hjz; *Blue*) and Apo-GLIC (*Yellow*) structures. The inter-SU pocket is either occupied by the CBX molecule (*A*, FUMAR, *B*, SUCCIN; *Yellow stick*), or empty (in the Apo-structure; *AB*). The intra-SU pocket is occupied by a chloride in the Apo-GLIC structure (*Green sphere*; *AB*), and also in the FUMAR-structure (*Light green sphere*, *A*), but occupied by the CBX molecule in the SUCCIN-structure (*B*). The Arg77 side-chain is oriented toward the intra-SU pocket in the Apo-structure (*AB*), and toward the inter-SU pocket in both CBX-structures (*AB*), showing a major impact of inter-SU pocket occupancy (*vs* intra-SU occupancy) on Arg77 side-chain orientation. A sodium is represented (*Purple sphere* in Apo-structure, *Dark pink sphere* in FUMAR- and SUCCIN-structures). A second chloride, unmodified between Apo- and CBX-structures, is visible in *A* and *B* (*Dark green sphere*). PDB references from Fourati *et al*. 2020. *C*, Data obtained using the *Xenopus* oocyte expression system. Bar graphs showing for WT GLIC and single mutants the values of the pHo corresponding to half maximal activation by extracellular pH (pHo_50_, *Left*), and the difference between mutant and WT GLIC pHo_50_ values (either n=1, or mean of 2 WT, obtained from the same injection) (ΔpHo_50_, *Right*). Number of cells tested (from at least two injections) indicated in brackets. Stars (*) left to two mutant names indicate that previously published data (Nemecz *et al*. 2017) were included. Pink and green colored areas indicate the mutant inclusion criteria: ± 0.5 pH unit from the WT pHo_50_ value. Y23A was excluded from the residue-dependency study. In this representation, data for a gain of function variant goes right to WT.

Each amino-acid was replaced by an Ala. The resulting GLIC constructs were characterized both in tk-ts13 cells (1-3 cells each), and in *Xenopus* oocyte recording (3-6 cells each, from at least two injections), for any shift in their proton sensitivity, evaluated for each cell as a pHo_50_ value, and a ΔpHo_50_ in comparison with a curve obtained from WT GLIC (1-2 cells) on the same day. Whole-cell patch-clamp and oocyte voltage-clamp gave consistent results, and oocyte data are presented in Fig. 6*C*. Gain of function mutants (decreased EC_50_) correspond to a decreased proton activity (a_H+o_) at half maximal current, *i*.*e*. an increased pHo_50_, and positive ΔpHo_50_. All mutants were found to give a rise in cation conductance during stimulation with low pH extracellular solution. E104A, which gives small GLIC currents and a slightly decreased pHo_50_ (see Fig. 2 in Nemecz *et al*. 2017), was replaced by E104Q. The Y23A variant showed a strong loss of function property, with almost one pH-unit decrease in pHo_50_ in both tk-ts13 (n=3) and oocyte (n=4) recording. The test pHo of 5.0 was near threshold for GLIC Y23A activation, so that the 8-10-fold CBX-to-control current *ratio* (10^3^ %) observed for SUCCIN or FUMAR at pHo 5.0 on Y23A was thought to be due to the loss of function property of the Y23A variant, and not further interpreted, except that Y23A does not suppress the PAM effect. The N152A variant showed a weak loss of function property, at the limit of inclusion, and also produced increased FUMAR PAM effects. The conclusion that N152A does not suppress the di-CBX PAM effect was unexpected, as Asn152 is in contact with the second carboxyl group in the GLIC-di-CBX cocrystal structures. The di-CBX and CAFFE data for N152A is presented and discussed with the mono-CBX data in the accompanying report (Van Renterghem *et al*. Manuscript M2), as a consistent explanation appeared in comparing mono-CBX and di-CBX N152A data. All other constructs presented a pHo_50_ within the inclusion *criterium* (± 0.5 pH-unit from the WT value; *i*.*e*. *ratio* of EC_50_ values: 0.3 to 3), and were used for this study.

**Figure 7.**
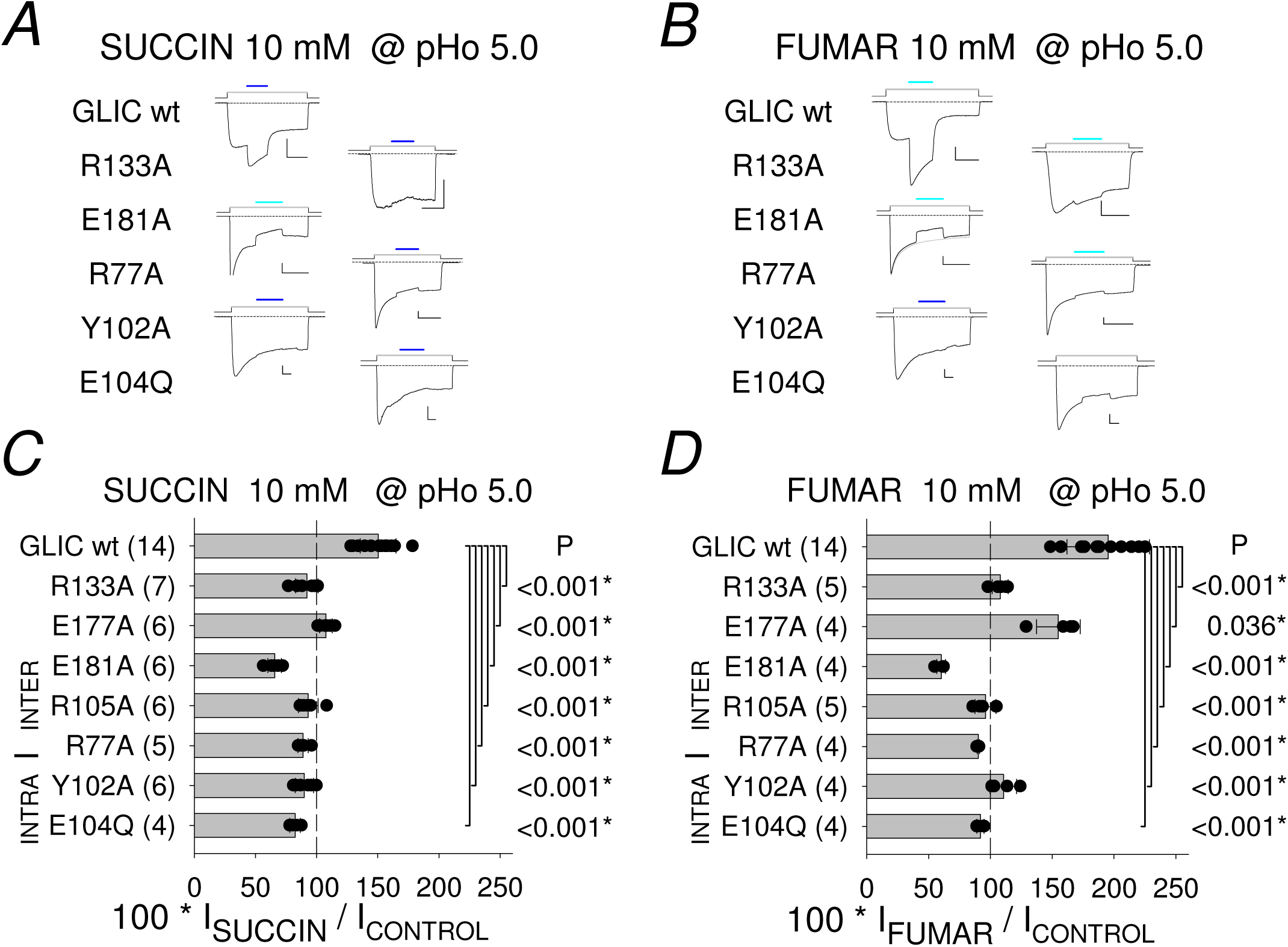
CBX-pockets single mutations: impact on di-CBX PM-effects. A, B, Representative current traces obtained for SUCCIN (*A*) and FUMAR (*B*) tests (10 mM at pHo 5.0) on WT GLIC and single mutation GLIC variants. Scale bars: 0.4 nA, 20 s. C, D, Bar graphs of current values (in % of control pHo 5.0-elicited current) in the presence of the PAMs SUCCIN (*C*) and FUMAR (*D*), on WT and single-mutation GLIC variants. Residue-belonging to the orthotopic site/inter-SU pocket, or to the intra-SU pocket, is indicated, as well as the border/pivot Arg77 (*Line*). Di-CBXs (10 mM) were tested at pHo 5.0 in the pre-stimulation protocol. Individual data points are super-imposed to bars indicating mean ± SD. The number of cells tested is indicated in brackets. Statistics: Each sample of values for a mutant was compared to the corresponding sample for WT GLIC, and Student’s T-test P values are indicated right to the bars.

#### Residue dependency for FUMAR and SUCCIN PAM effects

FUMAR and SUCCIN were tested at a concentration (10 mM) resulting in the maximum potentiation observed at pHo 5.0 on GLIC WT (see Fig. 1*B*). The single mutations in the inter-SU CBX-binding pocket (R77A, R105A, E181A) and its peripheral entrance (R133A, E177A) abolished the PAM effect (Fig. 7), except that E177A impact was only a reduction in the test with FUMAR (Fig. 7*D*). Surprisingly, the two single mutations in the intra-SU pocket (Y102A, E104Q) also fully suppressed the PAM effects (Fig. 7). In addition, E181A converted the di-CBX compounds into inhibitors (Fig. 7).

From this unexpected “all-or-none” pattern of residue dependency in the two binding pockets plus the orthotopic entrance, we conclude that the di-CBX PAM effect behaves as an “AND” system, requiring the integrity of both regions corresponding to the inter-SU pocket / orthotopic site, and to the intra-SU (= vestibular) pocket.

**Figure 8.**
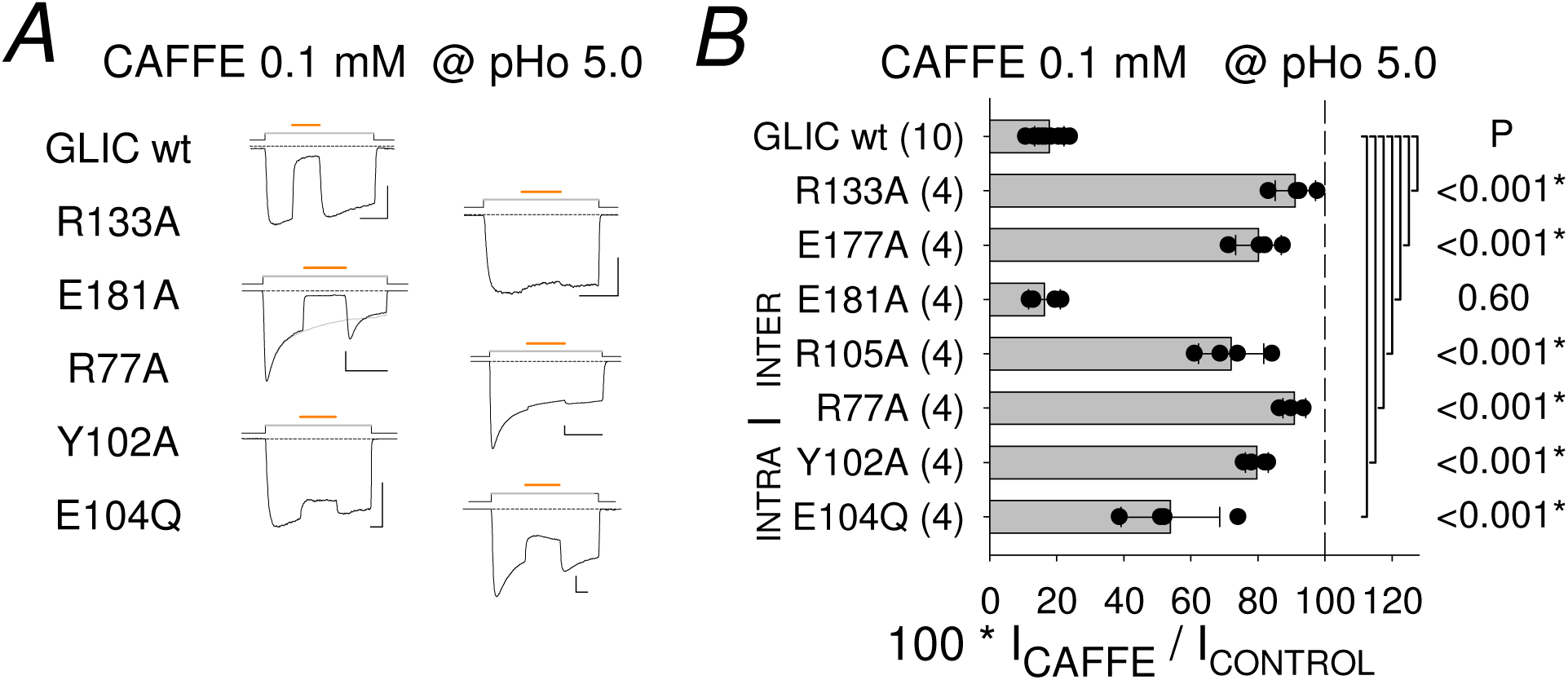
CBX-pockets single mutations: impact on CAFFE NM-effect. Representative current traces (*A*) and bar graph (*B*) for CAFFE tests (0.1 mM at pHo 5) on WT GLIC and single mutation GLIC variants. Scale bars: 0.4 nA, 20 s.

#### Residue dependency for caffeate inhibitory effect

Caffeate (CAFFE) was previously identified as a negative modulator of allosteric transitions (NAM) of GLIC, active at low concentration (IC_50_ = 17 µM), and a mutational analysis of the residue dependency at pHo 5.5 in oocytes concluded that CAFFE effect may occur through the orthosteric/orthotopic site of GLIC (Prevost *et al*. 2013). Tests with CAFFE performed in the present work on the same mutants, at the same, near IC_80_ CAFFE concentration (0.1 mM), but at pHo 5.0, produced similar results (Fig. 8): CAFFE inhibitory effect, leading on the WT to 18 (± 4, n = 10) % of the control current, was abolished or strongly reduced in R77A (91 %, ± 3 %, n = 4, P <0.001), R105A (72 %, ± 10 %, n = 4, P <0.001), R133A (91 %, ± 6 %, n = 4, P <0.001), and E177A (80 %, ± 7 %, n = 4, P <0.001), and not modified on E181A (16 %, ± 5 %, n = 4, P = 0.61).

In addition to previous works, we report that CAFFE inhibitory effect was also strongly reduced in mutants of the intra-SU pocket (Fig. 8), with 80 (± 4, n = 4, P <0.001) % of control on Y102A, and 54 (± 15, n = 4, P<0.001) % of control on E104Q, *vs* 18 (± 4, n = 10) % of control on WT as mentioned.

Therefore, the di-CBX PAM effect and the CAFFE NAM effect show a similar “all-or-none” pattern of residue dependency except in Loop C, since E181A does not suppress the CAFFE NAM effect, and E177A does not fully suppress the di-CBX PAM effect. We conclude here that the “AND” system applies to negative modulation and positive modulation, with the integrity of the orthotopic site/inter-SU site and the vestibular/intra-SU site both required for compound PAM and NAM effects.

## Disscussion

### Positive modulation of GLIC, with conditions of pHo

In this report, short-chain di-CBX compounds are identified as PAMs of GLIC (at pHo 5), with an optimal 4-carbon chain length and optimal constrained shape of the molecule, since FUMAR is the most potent compound identified, followed by the saturated 4-carbon di-CBX SUCCIN, whereas the *cis* compound MALE is inactive. This supports the view that the di-CBXs exert their potentiating action by binding through two pharmacophores at opposite ends of the molecule, which are expected to be the two carboxyl groups. The flexible SUCCIN molecule may need to adopt an anti-periplanar conformation to become a GLIC PM. The conformational flexibility of SUCCIN most likely lowers its potency, in comparison with FUMAR, which is already constrained in the optimal geometry. A clear selectivity appears towards other compounds, including aspartate and the neurotransmitter glutamate, which are inactive on GLIC.

According to the allosteric model (Monod e*t al*. 1965, Rubin & Changeux 1966) a PAM effect is expected to be larger in the lower part of the agonist activation curve. The di-CBX PAM effect, however, occurs only in a paradoxical low-pHo window (4-5), *i*.*e*. in the upper part of the GLIC activation curve (>EC_50_), thus determining the absence of agonist effect at pHo 7.5. Protonation/deprotonation of the di-carboxylic acid molecules occurs precisely in the range of pH (6-4) required for GLIC threshold to maximum activation (thought to depend on protonation/deprotonation of carboxyl groups in the side-chains of Asp and Glu residues, Nemecz *et al*. 2017). Whether this pHo-window for a PAM effect results from the influence of pH on the compound species distribution (see Table 1), or/ and on the GLIC protein, is therefore difficult to estimate. Regarding the pentamer, (de)protonation of residues belonging to the di-CBX binding site(s) may influence the ability of the site(s) to bind the compounds. Side-chains (de)protonation may occur as well along the binding-to-gating transmission pathway(s), conditioning transmission properties. (De)protonation in the protein would then determine the positive *vs* negative influence of a compound, in addition to directly controlling the probability for various receptor-channel conformations.

In search of a compound with agonist properties on GLIC, we found PAMs which are active only at low pHo. The question arises whether no agonist at all (at pHo near 7.5) will ever be found for GLIC, due to a question of permissive pHo for pore gating and/or permeation.

### Open state recruited with FUMAR@pHo 5 and with low-pHo only

Potentiation of I(pHo 5) by FUMAR occurred with no obvious change in ion selectivity (as judged from P_Cs_/P_Na_), suggesting that a common conformation of the pore is recruited as the open state. This suggests that the selective binding of a compound to a specific site (thought to be the orthotopic agonist site) does not further alter the open-pore conformation which results from the influence of pHo (a physical parameter) at numerous locations in the pentamer. GLIC is known as a (“non-selective”) cation channel (P_Cl_/P_Na_ = 0), with a sodium to potassium ions permeability *ratio* of 1.3 ± 0.2 (P_K_/P_Na_= 0.77), independent of pHo (Bocquet *et al*. 2007). The low permeability of caesium ions reported here (P_Cs_/P_Na_= 0.54) shows that GLIC ion pore discriminates between caesium and sodium ions, whether FUMAR is present or not. This observation, and the known small single channel conductance (8-9 pS), are consistent with the small diameter of GLIC ion pore observed in the crystal structures (without and with FUMAR), and support the interpretation that the structures obtained at pH 4 actually correspond to the open state of GLIC.

### The inter-SU pocket /orthotopic site and the intra-SU (= vestibular) pocket are functionally interdependent. A model with orthotopic site ligand efficiency conditioned by the integrity of the vestibular region

It is now clear that it is not in the orthotopic site, but in other locations, that protonation/deprotonation of residues determines the pHo-dependent control of GLIC gating (Nemecz *et al*. 2017). Showing the functionality, especially in positive modulation of channel activity, of the orthotopic site in the GLIC ancestor, which has been used as a pLGIC model in structural studies, is a major point, since this region, homologous to the neurotransmitter binding site, is the reference agonist site for pLGICs.

Analysis of the residue-dependency for the di-CBX PAM effect reveals the necessary integrity of two distinct regions in the GLIC ECD. The required orthotopic site region covers the deeply buried inter-SU CBX-binding pocket, as well as its slightly more peripheral entrance belonging to the pLGIC orthotopic site. The intra-SU CBX-binding pocket in GLIC coincides exactly with the vestibular pocket identified in ELIC. The “all-or-none” impact in the two pockets indicates that potentiation by the di-CBX compounds acts as an “AND” system, *i*.e. requires the simultaneous integrity of both sites

Two hypotheses may be derived from this observation (Fig. 9). The first one is that binding at both sites simultaneously is required for an effect (double occupancy). The second hypothesis is that binding occurs at only one of the sites (single occupancy), and the other pocket is involved functionally (see below). In the context of a single occupancy, one hypothesis (among two) is that binding occurs only at the inter-SU pocket, as neurotransmitters bind to the orthotopic site in Eukaryote receptors. Involvement of the inter-SU pocket as the major di-CBX binding site is supported by crystallographic data from GLIC-di-CBX co-crystals, showing that FUMAR and SUCCIN are both found occupying the inter-SU pocket in their respective co-crystals (Fourati *et al*. 2020). The region corresponding to the vestibular pocket would then condition inter-SU/orthotopic binding itself, or/and the transmission of forces between the orthotopic site and the rest of the protein during allosteric transitions, determining the gating probabilities.

**Figure 9.**
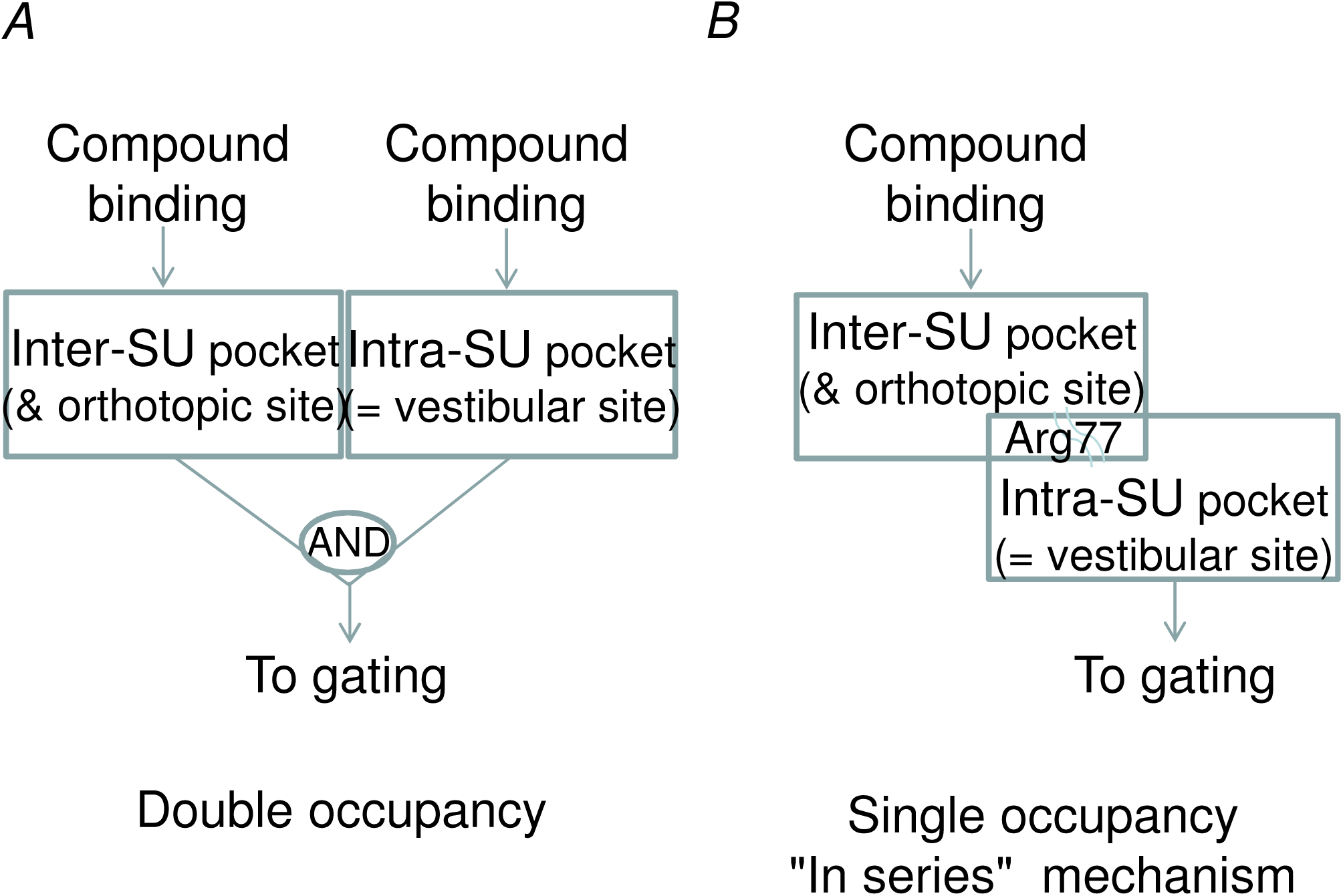
A single-occupancy mechanism with vestibular control may lead to a functionally 2 site-dependent system. Two hypothesis compatible with the “all or none” pattern of residue-dependency in the two CBX-binding pockets derived from single mutation data. *A*, Double occupancy mechanism, and *B*, a single occupancy model, with two sites “in series”, and vestibular control of the coupling between orthotopic/inter-SU ligand binding and gating.

Given that the same pattern of residue-dependency occurs for CAFFE inhibitory effect (with the exception of E181A, in Loop C), this speculation may be extended with the hypothesis that CAFFE as well binds at the inter-SU/orthotopic site, as previously proposed by Prevost *et al*. (2013). Binding to the inter-SU site would then promote positive (di-CBXs) or negative (CAFFE) modulation, depending on compounds, and depending on additional protein determinants. In a homology with the mechanism of benzodiazepines and beta-carbolines modulation of GABA-activated GABA_A_ receptors, the di-CBXs and CAFFE would then function respectively as positive and negative modulators of the low-pHo activated GLIC. In other words, positive and negative modulations occur due to the (alternative) binding of distinct compounds to the same (orthotopic) binding site, or strongly overlapping inter-SU binding site.

Our mutational analysis identifies one residue differentially involved in positive and negative modulation. Indeed Glu181, common to the orthotopic site (Loop C) and the inter-SU CBX binding pocket, is required for di-CBX potentiation (Fig. 7), but not required for CAFFE inhibition (Fig. 8). The fact that E181A (not E177A) converts FUMAR and SUCCIN into negative modulators further supports a differential requirement of Glu181 for positive, not negative modulation.

### Previously published crystallographic data support this model in GLIC

Interaction between inter- and intra-SU pockets in GLIC is supported by crystallographic data regarding Arg77, the amino-acid located at the interface of the two pockets (Sauguet *et al*. 2013, Fourati *et al*. 2015, 2020). As already noted by these authors, Arg77 handles an inter-SU CBX-associated pivot movement from the vestibular pocket to the inter-SU pocket (Fig. 6*AB*). Indeed, Arg77 side chain is oriented towards the vestibular pocket (occupied by a chloride ion) in the apo-GLIC structure obtained using phosphate as pH buffer (PDB reference 4qh5; Fourati *et al*. 2015). Whereas, in all the CBX-bound GLIC structures available, it is found coordinating the CBX-molecule bound into the inter-SU pocket, whatever is the *status* of the vestibular pocket (occupied by a CBX molecule or occupied by a chloride ion/’CBX-empty’) (Sauguet *et al*. 2013, Fourati *et al*. 2020). This includes four structures of GLIC in complex with molecules of the di-CBX compounds analyzed in the present study, malonate (6hjb), succinate (6hjz), glutarate (6hja) and fumarate (6hj3). We display in Fig. 6*A* the Arg77 side-chain pivot movement in the case of FUMAR (6hj3) *vs* apo-GLIC (4qh5). We also display the positions of Arg77 side chain in the SUCCIN-bound GLIC structure (6hjz) *vs* apo-GLIC structure (Fig. 6*B*). Arg77 points to the inter-SU pocket, occupied by FUMAR or SUCCIN respectively, despite the fact that a SUCCIN molecule has been assigned to the intra-SU pocket as well in 6hjz. It may be noted here that inter-SU pocket occupancy seems to be more robust than intra-SU pocket occupancy, as judged considering the 6hj3- and 6hjz-associated electron densities deposited by Fourati *et al*. (2020). The five inter-SU pockets of the pentamer are clearly occupied by FUMAR or SUCCIN molecules respectively. But regarding the intra-SU pockets, only one of the ten (with two pentamers in the crystallographic asymmetric unit) 6hjz-associated densities allows for a convincing placement of a SUCCIN molecule, suggesting that binding to the intra-SU pocket is weaker. FUMAR is clearly absent from the intra-SU pocket in 6hj3 densities (with densities corresponding to chloride atoms/ions), suggesting that the intra-SU pocket cannot handle the constrained shape of the FUMAR molecule. In any case, the inter-SU occupancy-associated Arg77 pivot movement, and the fact that inter-SU occupancy is observed in all the CBX-bound structures, both support the hypothesis that the orthotopic/interSU site is the major binding-site involved in the CBX pharmacological effects on GLIC.

In the single occupancy model with vestibular control, the two sites are “in series”, and compound binding at the most peripheral (orthotopic/inter-SU) site is sufficient to promote a PAM effect *if* the vestibular region is intact. As an extension of this scheme, it is conceivable that an additional vestibular binding may interfere in the resulting modulation, at least in this Prokaryote pLGIC. Vestibular binding would then either amplify or reduce the orthotopic-ligand elicited modulation of allosteric transitions. This scheme, with a principal inter-SU binding site, and a secondary binding site in the vestibular region, may be valid for several Prokaryote pLGICs, such as GLIC and may be ELIC. The question occurs whether vestibular binding may be the main (or single) binding component in other Prokaryote receptors, as proposed in the case of the sTeLIC (Hu *et al*. 2018).

### Relevance of the model to Eukaryote pLGICs

Evolution toward nervous system receptors has clearly favored the orthotopic site as the binding site for neurotransmitters, *i*.*e.* the binding site for the physiological agonist PAM effect. Binding to orthotopic sites leads to negative modulation in a few cases, through accessory orthotopic sites in heteromeric pLGICs, as with beta-carbolines on GABA_A_ receptors. And in peculiar cases, orthotopic antagonists, acting by binding competition with the orthotopic agonist, may reveal NAM properties on constitutively active variants. But involvement of the orthotopic site as the main, agonist binding site in Eukaryote pLGICs is obviously well established and has been extensively studied.

It appears of great interest to question in human pLGICs the putative involvement of the region homologous to the prokaryotic vestibular pocket in mediating the modulation driven by orthotopic agonist binding, as well as the ability of the vestibular region to become the binding target of new therapeutic modulators. Does the vestibular site exist in Eukaryote pLGICs? Is it functional? Is Arg77 conserved?

According to a multi-alignment of pLGICs amino-acid sequences, the extremely conserved motif WxPD/E preceding Arg77 (WIPE-IR_77_FVN) in GLIC is followed in most anionic pLGICs by three aromatic residues, and in most of nicotinic and 5HT3 receptors by three non-aromatic hydrophobic residues, while, two positions later, an extremely conserved Asn occurs. GLIC Arg77, in-between highly conserved residues, is therefore not conserved in the sequence of Eukaryote pLGICs. And the very simple and impressive lever arm coupling inter-SU and intra-SU pockets in GLIC was probably replaced during evolution by more complex, and may be more efficient, molecular components of orthotopic site/vestibular site coupling.

Fourati *et al*. (2015) already noted that in GluCl (3rif; deposited by Hibbs & Gouaux (2011), as in type β3 GABA_A_ receptor (4cof, Miller & Aricescu, 2014), the site occupied by an insertion of the pre-β5 loop (later called Loop Ω) of the neighboring sub-unit is exactly homologous to the vestibular pocket. In anionic pLGICs, the occupied vestibular pocket is therefore not accessible to compounds, or not accessible from the vestibule lumen, as confirmed with later works involving more pLGICs structures (Brams *et al*. 2020). However, Fourati *et al*. (2015) also noted that, in the alpha 1 type glycine receptor (α1GlyR), the position of Zn^2+^ in a model for the Zn^2+^ inhibition site (Miller *et al*. 2008) corresponds exactly to the position of the intra-SU bound acetate in GLIC. Mutations in this site abolish inhibition by Zn^2+^ (Miller *et al*. 2008). The vestibular site therefore seems to be functional as an allotopic binding site for the NAM effect of the divalent cation in human α1GlyRs. Regarding cationic pLGICs, Hu *et al*. (2018) noted that a vestibular cavity is open and accessible in the mouse 5HT_3A_ receptor. A systematic analysis of the vestibule site architecture and accessibility in all available prokaryote and eukaryote pLGIC structures was published in 2020 by Brams *et al*. Their analysis actually considers the luminal entrance of the vestibular pocket, and shows that the entrance is open in muscular type nicotinic subunits and 5HT_3A_ subunits. In addition, these authors show that MTSEA-biotin modification of engineered Cys mutants in the vestibule site (in particular Leu151 in the β6 strand) produces a PAM effect on the serotonin-activated 5HT_3A_ receptor, showing that it is possible to target the vestibule site for modulation of the 5HT_3A_ receptor, and putatively other eukaryote pLGICs.

### Influence of intracellular pH, GLIC current decay and desensitization

Although not providing a systematic study of GLIC pHi dependency, when either low-pHo only or [FUMAR at low pHo] appears as an extracellular stimulus, our whole-cell protocol data demonstrates that 1/ low-pHo induced GLIC current decreases when pHi decreases, and 2/ [FUMAR at pHo 5]-induced current decreases much less than pHo 5-induced current, so that, with tests performed at pHo 5, the FUMAR-induced potentiation *ratio* strongly increases when pHi decreases.

In our data with and without FUMAR, the cell and the pentamer are exposed to the same test-pHo value. The pHo 5-induced GLIC current is reduced with intracellular pH 6 to about one-fifth of its value with intracellular pH 7.3. The fact that the FUMAR-induced current is not reduced in the same proportion suggests that the inhibitory effect of low pHi is not (or not only) an effect on the permeation properties of the open pore, but an effect of pHi on the open state probability. In other words, the differential effect of pHi on data with and without FUMAR demonstrates that pHi modulates the allosteric transition (between conformational states), as does pHo.

Therefore, GLIC sensitivity to pHi is opposite, and as much a determinant as its sensitivity to pHo. This observation raises several theoretical points regarding GLIC functional properties. One question is whether GLIC is activated depending on absolute pH values inside and outside, or whether GLIC activation level depends on the transmembrane proton gradient, the pHo – pHi difference, or a_H+_o/a _H+_i proton activity *ratio*.

Another question raised is how much a progressive drop in pHi, consequent to the experimental lowering of pHo, contributes to GLIC current decay, in electrophysiological studies. As reported by other authors (Laha e*t al*. 2013), we commonly observe a great variability in the kinetics of decay of the current recorded at a given pHo value, particularly in conditions with poor intracellular solution control, hence poor buffering (incomplete cell dialysis at short whole-cell recording time or at high series resistance intracellular access). A low-pHo induced progressive drop in sub-membrane intracellular pH may be a major determinant of GLIC current decay.

The question whether a proper desensitized (D) state exists in GLIC or not is complicated by the fact that a dropping-pHi induced current decay may be (mis)-interpreted as desensitization. Questioning a D state in GLIC leads to questioning the definition of the D state. Would it be favored and recruited after a long exposure at low-pHo? Would it be recruited after a long exposure to an (unknown) agonist, or a PAM such as FUMAR, and would this occur only if this compound is an orthotopic-site ligand?

In Eukaryote receptors, criteria for a D state include a closed channel state, with high affinity for the agonist. In electrophysiology, with macroscopic data, an agonist-induced transition to a D state is observed as current decay in the presence of the agonist. The channel can leave the D state and go back to a resting (R) state after agonist washout. Kinetics of the transitions to and from the D state are usually slower than transitions to and from the active, open state, allowing to observe desensitization from the current decay, and to follow de-desensitization using brief agonist tests.

Extracellular GLIC activation/potentiation, always associated with experimentally imposed low–pHo (putatively promoting proton influx through GLIC ion pore and transporters), will always promote a low–pHi situation, but indirectly, in a time and agonist/PAM concentration-dependent way. Recovery from low–pHo induced pHi–lowering is also expected to take time (more or less depending on the pHi level reached, and depending on numerous parameters, such as membrane transporters, ion concentrations, cellular and pipette-provided pH buffers, *etc*.) So that dropping–pHi effects in whole-cell recording behave very much like a proper desensitization.

One hypothesis may reconcile GLIC properties with the concept of desensitization: low–pHi conditions would stabilize a desensitized state, defined on a functional basis, as non-activatable by an extracellular stimulus. But data with FUMAR when pHi is very low would be required to examine this point.

It may be that a low–pHo induced, dropping–pHi (rising-H^+^)–mediated channel inactivation appeared useful to a cell’s survival, and that evolution developed the proper agonist-compound-induced desensitization as a process which mimics a useful agonist H^+^_o_-induced, H^+^_i_-mediated inactivation process. A similar evolutionary concept regarding voltage-gated channels may hypothesize that proper intrinsic voltage-dependent inactivation derives from the ancestral process of a rising–Ca^2+^_i_-mediated, calcium-influx-associated inactivation as observed nowadays in L-type calcium channels. However, whether a proper desensitization, elicited within the pentamer by extracellular compound binding, occurs or not in GLIC, in addition to the H^+^ -mediated process described here, the point remains open.

### Succinate and fumarate as ancestors of neurotransmitters?

Noticeably, a chemical proximity is obvious between the di-CBX compounds active on GLIC as PAMs, and the major excitatory neurotransmitter or related agonist, since glutamate and aspartate differ only by an alpha-amino group from GLUTAR and SUCCIN. *Gloeobacter violaceus*, a cyanobacterium lacking thylakoids (Rippka *et al*. 1974), is thought to be a surviving species of an extremely archaic period of evolution of the living world, with characteristics thought to correspond to Precambrian species. It may be that pLGICs appeared as “proton-gated” (or “pH-gated”) channels, and that organic compounds first appeared as modulators, and later evolved to become agonists and neurotransmitters (with usually unnoticed background conditions of pHo and pHi for channel opening and/or permeation), while the agonist effect of changes in pH was lost with the progressive establishment of a constant *milieu intérieur*. If it is the case, it may be expected that pH-dependent gating, occurring either way (as occurring in opposite ways for GLIC and sTeLIC), may be common among bacterial pLGICs.

### Kreb’s cycle components as positive modulators of a proton-activated cation channel: the “millstone round the neck” of *Gloeobacter violaceus*?

Another remarkable property of di-CBX compounds is their involvement in the citric acid cycle. GLIC-potentiating compounds (SUCCIN, FUMAR, and OAA) are involved in the second half of the Krebs’ cycle, whereas the entry compound, citrate, resulting from the combination of a new pyruvate (*via* acetyl-CoA) with OAA, is found to be inactive on GLIC. The location of GLIC in the *Gloeobacter violaceus* cyanobacterial cell has not been described, but the receptor-channel is clearly expected to be expressed in the plasma membrane and have its ECD exposed to the periplasmic space. The Krebs’s cycle metabolic pathway is likely located in the cytosol, as in other bacteria, and it is unclear whether Krebs’s cycle components or other CBX molecules can pop-up in the periplasmic space. This compartment is however likely to be a low-pH compartment, due to the function of both respiratory and photosynthetic systems in the plasma membrane (in the absence of thylakoids). It may be that, stimulated by energy-producing processes, GLIC would short-circuit cation gradients, and possibly the proton-gradient necessary for ATP production (especially if protons are permeant, as are other cations, a point which is probable but not demonstrated). The GLIC protein would then be labeled as the “millstone round the neck” of this species, known for its very long generation time (Rippka *et al*. 1974, Nakamura *et al*. 2003), a disadvantage for evolution, but an advantage for the durability of a peculiar species. If this hypothesis is correct, a knock-out of GLIC is expected to produce a faster-growing variant of *Gloeobacter violaceus* (which might endanger the species durability if dispersed in its natural environment).

## Abbreviations list

ASIC: acid-sensing ion channel
BAPTA: 1,2-bis(*o*-aminophenoxy)ethane-*N*,*N*,*N’*,*N’*-tetraacetic acid
BHK: baby hamster kidney
CAFFE: caffeic acid/caffeate
CBX: carboxylic acid/carboxylate
DCM: dichloromethane
dFBS: heat-inactivated fetal bovine serum
ECD: extracellular domain
ESI: electrospray ionization
FUMAR: fumaric acid/fumarate^-^/fumarate^2-^
GABA: *gamma*-aminobutyric acid
β3GABAAR: β3 type A GABA receptor
GLIC: *Gloeobacter violaceus* ligand-gated ion channel
α1GlyR: alpha 1 glycine receptor
HEPES: 4-(2-hydroxyethyl)-1-piperazine ethanesulfonic acid
inter-SU: inter-subunit
HPLC: high performance liquid chromatography
HRMS: high resolution mass spectrometry
Imax: active current extremum value (with largest absolute value) in a concentration-effect plot
Ipk: peak current active current extremum value (with largest absolute value) in a segment of recording time
MALE: maleic acid/maleate^-^/maleate^2-^
MES: 2-(*N*-morpholino) ethanesulfonic acid
mM: mmol/L
NAM: negative modulator of the allosteric transitions
NMDG: *N*-methyl-D-glucamine
NMR: nuclear magnetic resonance
OAA: oxaloacetic acid/oxaloacetate^-^/oxaloacetate^2-^
PDB: Protein Data Bank
PAM: positive modulator of the allosteric transitions
pHi: pH of solution at the intracellular face of the cell membrane
pHo: pH of solution at the extracellular face of the cell membrane
pHo_50_: extracellular solution pH value producing half of the maximal activation reached using low extracellular pH
pHpip: pH of the solution used to fill the recording pipette
pLGIC: pentameric ligand-gated ion channel
SUCCIN: succinic acid^-^/succinate^-^/succinate^2-^
SU: subunit
TFA: trifluoroacetic acid.

## Additional information

### Data Availability Statement

For patch-clamp electrophysiology data, representative raw current traces (current recorded *versus* time) are available in all Figures (Figs. 1-5, 7, 8). For electrophysiology data presented as bar graphs, individual value data points (one per cell in each condition) are superimposed to mean ± SD (Figs. 3, 4*B*, 5*CD*, 6*BC*, 7, 8), and the number of cells tested is indicated in brackets in each bar graph category name. Mean ± SD is represented in Graphs for [concentration] – [current] “dose-response” curves (Fig. 1*B*) and [relative current] – [time spend in whole-cell recording configuration] kinetic graphs (Fig. 3*BDF*). The number of cells included is then indicated in the Figure Legend. In all cases, “n” includes cells from at least two DNA transfections (tk-ts13 cells) or two DNA injections (oocytes). We provide in the present report a very detailed description of the methods used. A statistical comparison of pairs of samples was performed in the mutational analysis (Figs. 7*C*,*D*; 8*B*), and also when relevant for other data presented as bar graphs (Figs. 1*D*, 2*BD*, 4*B*, 5*F*). P values are indicated near bar graphs in the Figures, and most of the time also in Text.

### Competing interests

The authors declare no competing interest.

### Author contributions

CVR set up patch-clamp, designed and performed patch-clamp electrophysiology experiments (and host cells handling), analyzed data (including statistical treatment), and prepared patch-clamp Figures. ÁN performed *Xenopus* oocyte electrophysiology recordings and analysis (and oocyte injections) and prepared Fig. 6. SDC and DJ designed and performed bioisosteres choice and chemical synthesis. P-JC initiated and supervised the work. CVR wrote the first version of the article. All authors contributed to revising the article.

All authors have approved the final version of the manuscript and agree to be accountable for all aspects of the work. All persons designated as authors qualify for authorship, and all those who qualify for authorship are listed.

### Funding

ÁN was supported by the Agence nationale de la recherche (ANR grant 13-BSV8-0020 “Pentagate”). CVR and P-JC have permanent CNRS positions, SDC and DJ have permanent Paris-Saclay University positions. P-JC’s laboratory belongs to IP and CNRS. ANR, CNRS, IP and UP-S are supported by French nation public funding. IP is also supported by multiple individual donators.

## Ackowledgements

CNRS, MESRI, Université Paris-Saclay and ANR (ANR-13-BSV8-0020) are gratefully acknowledged for financial supports. The authors wish to thank Nicolas Bocquet, Virginie Dufresne, Zaineb Fourati, Christèle Huon, and Karima Medjebeur for some molecular biology constructs and DNA production, as well as Chantal Jan, Colette Jean and Corinne Balayira for technical assistance with lab-ware. They thank Karine Leblanc for HPLC and HRMS analyses, Laurie Peverini for two ^1^H NMR spectra at the end of patch-clamp experiments, Nicolas Lenovère for formerly providing a multi-alignment of pLGICs amino-acid sequences, and Jean-Marie Biansan for the free software Dozzzaqueux. CVR thanks Frédéric Poitevin, Ludovic Sauguet and Marc Delarue for useful discussions at the beginning of this work.

## First author Profile

After studies at Ecole Sainte-Geneviève (Versailles), Ecole normale supérieure de Fontenay-aux-Roses (now ENS Lyon), and Université de Paris (Jussieu), CVR chose electrophysiology after listening to lectures by Philippe Ascher. She did a PhD (1984) with Jacques Stinnakre in the laboratory headed by Ladislav Tauc (Gif-sur-Yvette). Then she joined the Group of Michel Lazdunski (Nice), and received a CNRS position. She chose to remain recording at the patch-clamp set-up, and worked in several laboratories on various topics (metabolic control of ion channels, voltage-gated Na and Ca channels, pre-synaptic toxins). She is now contributing to electrophysiology in the Group headed by Pierre-Jean Corringer.

**Authors’ Translational Perspective**

## Notes

### Competing Interest Statement

The authors have declared no competing interest.

